# The TUTase URT1 regulates the transcriptome of seeds and their primary dormancy

**DOI:** 10.1101/2024.12.19.629392

**Authors:** Jackson Peter, Jeanne Roignant, Sebastian Sacharowski, Elodie Ubrig, Benjamin Lefèvre, Szymon Swiezewski, Dominique Gagliardi, Hélène Zuber

## Abstract

RNA uridylation is a pervasive mechanism that regulates the degradation of eukaryotic mRNAs. In Arabidopsis, uridylation influences mRNA decay both by favoring 5’ to 3’ degradation and by preventing excessive deadenylation. Yet, the significance of mRNA uridylation during plant development remains largely unknown. Here, we adapted FLEP-seq2, a method based on nanopore sequencing, to generate a comprehensive inventory of mRNA uridylation events in different Arabidopsis tissues. We also evaluated the respective contribution of the two known Arabidopsis uridylyltransferases, URT1 and HESO1, in mRNA uridylation. Our transcriptome-wide analysis showed that, URT1 is the main enzyme responsible for mRNA uridylation, in all analyzed tissues, while HESO1 can marginally uridylate mRNAs. Importantly, our results revealed the singularity of mRNA uridylation pattern in seeds and the dual function of URT1-dependent uridylation in shaping the transcriptome during seed maturation. We propose that during the late stages of seed maturation, URT1-dependent uridylation facilitates the degradation of unnecessary mRNAs encoding translation-related proteins, while also promoting the accumulation of mRNAs associated with the maturation program, by hindering their deadenylation. In line with its important function in shaping the seed transcriptome, our study also identifies URT1 as a novel regulator of seed dormancy. Overall, our study reveals the biological relevance of mRNA uridylation during the late stages of seed maturation.

## Introduction

mRNA uridylation is an abundant post-transcriptional modification that is conserved in most eukaryotes, from fission yeast to plants and humans (De Almeida et al. 2018; Warkocki et al. 2018; Zigáčková and Vaňáčová 2018; Yu and Kim 2020). The addition of uridines (U) to the 3’ end of mRNAs is catalyzed by Terminal UridylylTransferases (TUTases) such as TUT4/TUT7 in mammals (Lim et al. 2014; Thomas et al. 2015; Morgan et al. 2017; Chang et al. 2018) or Cid1 in fission yeast (Rissland and Norbury 2009; Grochowski et al. 2024). In Arabidopsis, UTP:RNA URIDYLYLTRANSFERASE 1 (URT1) is the main enzyme responsible for mRNA uridylation (Sement et al. 2013; Zuber et al. 2016). Yet, the residual uridylation detected in the *urt1* null mutation indicates that Arabidopsis contains at least a second enzyme able to uridylate mRNAs. HEN1

SUPPRESSOR1 (HESO1), the other prominent TUTase characterized in Arabidopsis, is involved in the uridylation of small RNAs and 5’-cleavage fragments resulting from cleavage by the RNA-induced silencing complex (RISC) (Ren et al. 2012, 2014, 2023; Zhao et al. 2012; Zuber et al. 2018; Vigh et al. 2024) and shared common RNA substrates with URT1 including some microRNAs (Tu et al. 2015; Wang et al. 2015), some 5’-cleavage fragments from RISC-cleavage (Zuber et al. 2018; Vigh et al. 2024) and the polyadenylated viral RNA of TuMV (Joly et al. 2023). While its role in mRNA uridylation has not been investigated yet, HESO1 is a good candidate as the alternative TUTase for mRNA uridylation. In eukaryotes, the main function of mRNA uridylation is to assist mRNA decay. In fission yeast or human cells, mRNA uridylation was described to facilitate the degradation of mRNAs from 5’ to 3’, by recruiting the decapping machinery and thus triggering the decay by 5′-3′ exoribonuclease 1 (XRN1), or alternatively from 3’ to 5’ by promoting the activity of Dis3-like protein 2 (Dis3L2) or the RNA exosome (Chowdhury et al. 2007; Song and Kiledjian 2007; Rissland and Norbury 2009; Malecki et al. 2013; Lim et al. 2014). In Arabidopsis, mRNA uridylation also participates in mRNA decay pathways. URT1 is part of a protein network comprising the deadenylation complex CCR4-NOT and translation repressors/decapping activators. Notably, URT1 directly interacts with DECAPPING 5 (DCP5), a protein that promotes mRNA decapping and participates in translation inhibition. Moreover, recent results based on direct RNA sequencing (DRS) using Oxford Nanopore Technologies (ONT) revealed a global role of URT1 in shaping the poly(A) tail length of mRNAs. URT1-mediated uridylation was shown to prevent the accumulation of excessively deadenylated mRNAs, likely by both favoring their 5’ to 3’ degradation and hindering excessive deadenylation. Importantly, this function protects plants against the biogenesis of illegitimate siRNAs that silence endogenous mRNAs and strongly impact plant growth and development (Scheer et al. 2021). Altogether, these results reflect the complexity of the molecular mechanisms associated with mRNA uridylation in plants and of its consequences on mRNA fate.

In mammals, mRNA uridylation has been shown to be essential for oocyte maturation and fertility (Morgan et al. 2017). In Arabidopsis, functional mRNA uridylation and degradation by the cytosolic RNA exosome are required to prevent that a subset of mRNAs associated with photosynthesis enters the siRNA pathway, highlighting the physiological significance of RNA uridylation (Wang et al. 2022). Yet, the regulatory roles of mRNA uridylation during a plant development process remains elusive and so far, knowledge about mRNA uridylation has mainly been collected in flowers and in seedlings. A comprehensive gene-to-gene analysis of mRNA uridylation profiles and of the contribution of URT1 and HESO1 in different plant tissues and stages is still lacking. FLEP-seq2, a Nanopore-based cDNA sequencing method, was recently developed to measure poly(A) tail sizes of mRNAs in a transcriptome-wide manner (Jia et al. 2022). This long-read sequencing method has the significant advantage to circumvent an inherent bias of Illumina sequencing toward poly(A) size estimation and to avoid purifying polyadenylated mRNAs. It is theoretically possible to use FLEP-seq2 to detect and analyze 3’ terminal added non-adenosine nucleotides and therefore uridylation. In the present study, we developed an additional bioinformatic module to analyze the composition and the length of mRNA non-adenosine tails and we showed that FLEP-seq2 can be used to accurately characterize U-tails. Using FLEP-seq2, we generated an atlas of mRNA U-tails in different plant tissues, including in developing, dry mature and germinating seeds for wild-type plants and *urt1* and *heso1* single or double mutants. The most striking observation was a strong accumulation of uridylated mRNAs, coupled to a lengthening of U-tails, specifically detected in seeds. Moreover, we showed that URT1 was the main contributor to this uridylation and that its loss has a prominent impact on the transcriptome of dry seeds and on seed primary dormancy. Finally, our findings revealed the importance of mRNA uridylation dual roles during a developmental process: during seed maturation, URT1-mediated uridylation promotes the downregulation of mRNAs encoding translation-related proteins, while facilitating the accumulation of mRNAs associated with the seed maturation program.

## Results

### mRNAs of dry mature seeds are highly uridylated

To first assess the accuracy of FLEP-seq2 to analyze uridines added to the RNA 3’ end, we sequenced synthetic DNA fragments harboring a 10-nt poly(A) tail followed by 0 to 6 thymidines (Ts). We improved the original version of the FLEP-seq2 analysis pipeline (Jia et al. 2022) by including an additional module to analyze the composition and the length of non-adenosine tails (https://github.com/jackson-peter/FLEPseq2). Our results showed that 91 to 96% of sequences corresponding to DNA fragments with one or several Ts were correctly tagged as uridylated RNAs while the technical background, *i.e.* percentage of sequences mistakenly tagged as uridylated RNAs, was lower than 1% (Supplementary Figure 1A, Supplementary Data Set S1). The number of Ts was more accurately estimated for shorter tails than for longer tails. For the synthetic DNA fragment containing 6 Ts, this number was correctly estimated for 59% of sequences and underestimated at 5 for 28% of sequences. Importantly, the number of Ts was accurately estimated (>90%) for synthetic DNA fragments containing 1 to 3 Ts (Supplementary Figure 1B, Supplementary Data Set S1). This accuracy is important because the vast majority of uridylated mRNAs detected in Arabidopis contain from 1 to 3 Us (Sement et al. 2013; Zuber et al. 2016). Altogether these results demonstrate good performances and accuracy of our pipeline for quantifying levels of uridylation and differentiating short and long U-tails from FLEP-seq2 data.

To perform a comprehensive analysis of mRNA U-tailing in different Arabidopsis plant tissues, we took advantage of FLEP-seq2 datasets from (Jia et al. 2022) and we analyzed uridylation profiles for nuclear-encoded mRNAs. Our results showed consistent overall percentages of uridylation across replicates ranging from 5 to 6% in most tissues (Figure 1A). Interestingly, the highest level of uridylation was found in dry seeds for which uridylation rate reached 9% on average. This represents an increase of 70% when compared to flowers. We then calculated the percentage of uridylation per mRNA and plotted the distribution of percentages for each replicate and tissue (hereafter refer to as the per-transcript distribution), where each transcript contributes equally to the distribution (Figure 1B, Supplementary Figure S1C). The per-transcript distribution showed higher uridylation rates in seeds regardless of the mRNA population, whether calculated for all mRNAs (Figure 1B), for those detected in all tissues or for those specifically expressed in dry seeds (Supplementary Figure S1C). Therefore, the overall high uridylation rate of mRNAs cannot be explained by the specificities of the seed transcriptome. Interestingly, we also observed a lengthening of U-tails in dry seeds. In all tissues except seeds, U-tails were mainly short, composed of 1 or 2 Us with less than 26% of tails longer than 2 Us (Figure 1C). This result confirms previous results obtained in flowers and in leaves using alternative methods (Sement et al. 2013). By contrast in dry seeds, the proportion of long U-tails substantially increased with 55% of U-tails longer than 2 Us. These findings highlight both an accumulation and a lengthening of U-tails in dry seeds as compared to other tissues.

**Figure 1:**
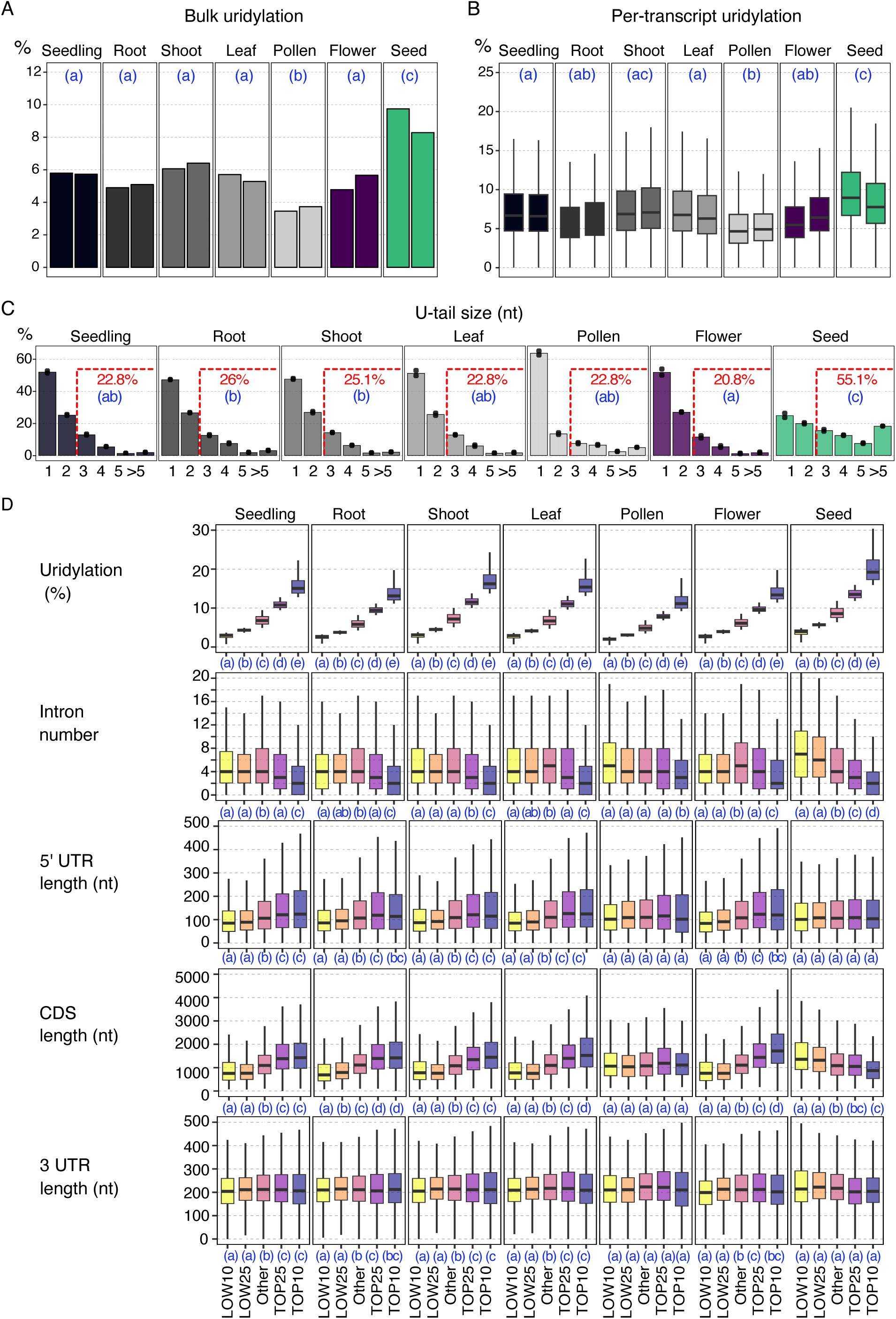
Comprehensive analysis of mRNA U-tailing in different Arabidopsis plant tissues. FLEPseq2 data from Jia et al. 2022 were re-analyzed with a pipeline adapted to analyze U-tails. Data are shown for two replicates. A-B, Bulk (A) and per-transcript (B) percentages of uridylation. Per-transcript percentages were calculated for mRNAs detected with at least 50 reads for each replicate. C, % of tails according to the number of Us. Individual points and the bar plot show individual replicates and the average, respectively. The average percentage of tail > 2 Us are indicated above each bar plot. In (A), (B), and (C), significantly different values are labelled by different letters (adjusted p-value < 0.05, n=3). A generalized linear model for proportion with a quasibinomial distribution was used to compared percentages in (A) and (C) and a linear model to compare medians in (B). D, Feature analysis for the 10% and 25% most and less uridylated mRNAs or for other mRNAs. Each boxplot displays the median, first and third quartiles (lower and upper hinges), the largest value within 1.5 times the interquartile range above the upper hinge (upper whisker) and the smallest value within 1.5 times the interquartile range below the lower hinge (lower whiskers). Significantly different values are labelled by different letters (adjusted p-value < 0.0001, unpaired and two-tailed pairwise Wilcoxon rank sum tests, n > 400).

### Uridylation preferentially targets mRNAs regulated during seed maturation

We then wondered whether some genomic characteristics were associated with a high or low mRNA uridylation rates and in particular whether some of these features could be specific to dry seeds. In dry seeds, highly uridylated mRNAs had both a significant lower number of introns and a shorter CDS length, while weakly uridylated mRNAs had a higher number of intron and a longer CDS length (Figure 1D). Intriguingly, the relationship between CDS length and uridylation is opposite in tissues other than seeds with highly uridylated mRNAs having longer CDS length in all other tissues except in pollen (Figure 1D). We then performed a gene ontology analysis and looked for biological processes significantly enriched among highly uridylated mRNAs. While in all tissues except pollen, gene ontologies associated with DNA transcription were significantly overrepresented among the 10% most uridylated mRNAs (hereafter called Top10% uridylated mRNAs), the biological processes associated with response to water deprivation, wounding and bacterium were specifically enriched in dry seeds (Supplementary Figure S1D and Supplementary Data S1). Of note, the most enriched gene ontology among Top10% uridylated mRNAs in seeds was the cellular response to hypoxia (False Discovery Rate (FDR): 1,92E-17, Fold Enrichment: 4.4) which was also significantly enriched but to a lesser extent in pollen (FDR: 0.02, Fold Enrichment: 3.7). Among the most uridylated mRNAs in dry seeds, we found many key genes of the seed maturation program, such as regulators of the abscisic acid pathway (HIGHLY ABA-INDUCED PP2C GENE 1, 2 and 3; ABI five-binding protein 1 and 2), the chlorophyll degradation (STAY-GREEN 1 and 2) or regulators of seed dormancy (DELAY OF GERMINATION 1 (DOG1); SLEEPY1). We expanded this gene ontology analysis to mRNAs, which were not necessarily the most uridylated but which showed the highest increase in uridylation levels in dry seeds when compared to other tissues. To identify these mRNAs, we plotted an heatmap showing for each mRNA the variation patterns of uridylation percentages between tissues (Supplementary Figure S1E). Out of the 1987 mRNAs detected with at least 50 reads in all tissues, the 869 mRNAs belonging to the cluster 1 showed the highest increase in the uridylation level in dry seeds compared to other tissues with an average 2.2-fold increase. Biological processes enriched for cluster 1 differed from those enriched among Top10% uridylated mRNAs and were associated with ribosome (FDR: 2.82-22, Fold Enrichment: 1.9) and translation (FDR: 8.32E-23, Fold Enrichment: 1.9, Supplementary Data Set S1). In line with this clustering analysis, the per-transcript distribution confirmed that the uridylation rate of most mRNAs associated with translation substantially increase in dry seeds compared to all other tissues (Supplementary Figure S1F). Interestingly, the end of seed maturation was previously associated with a gradual decrease in translation at the end of the seed maturation (Bai et al. 2023). Altogether, the FLEP-seq2 data revealed that uridylation preferentially targets major classes of mRNAs that are associated with important and highly regulated processes of the seed maturation.

### Distinct processes explain high uridylation rates in developing and dry seeds

In order to confirm our analysis based on results from (Jia et al. 2022) datasets and to obtain further information about the dynamic of uridylation profiles in seeds, we produced FLEP-seq2 libraries from three biological replicates for different seed stages: developing, dry mature and germinating seeds (Figure 2A). For developing seeds, we analyzed a pool of developing stages from embryogenesis to seed filling (∼ from 5 to 14 days after flowering, before the onset of desiccation). We analyzed both isolated seeds and siliques containing the seeds. For germinating seeds, we analyzed seeds imbibed in water for 24 hours. For comparison, we produced libraries from flowers. Our analysis first confirmed previous observations: both bulk and per-transcript percentages of uridylation were two-fold higher in dry seeds (bulk uridylation of 11% on average) than in flowers (bulk uridylation of 5% on average) (Figure 2B and C, Supplementary Data Set S2). This increased uridylation rate was not restricted to seed specific mRNAs (Supplementary Figure S2A), and U-tails were longer in dry seeds with on average 60% of U-tails containing more than 2 Us compared to 23% in flowers (Figure 2D). Moreover, as previously observed, highly uridylated mRNAs in dry seeds have a lower number of introns and a shorter CDS length (Supplementary Figure S2B). Interestingly, our new analysis also highlighted changes of mRNA uridylation rates and profiles during seed development and germination. In developing seeds or siliques, the mRNA uridylation rate reached a similar, albeit a little lower level (9.4-9.9%) as compared with the uridylation rate observed in dry seeds (Figure 2B and C). By contrast in germinating seeds, this rate decreased to 5% on average, similar to what was observed in flowers. This observation implies a highly dynamic regulation of mRNA uridylation rates during the first stages of seed germination. The length of U-tails was also modulated during development and germination with U-tails being longer in dry seeds compared to developing seeds (25% of U-tails longer than 2 Us in developing seeds) and shortened again in germinating seeds (only 16% of U-tails longer than 2 Us in developing seeds). Finally, the relationship between the CDS length and the uridylation level of mRNAs differs between dry seeds and developing or germinating seeds: CDS of highly uridylated mRNAs were significantly shorter in dry seeds but longer in both developing and germinating seeds (Supplementary Figure S2B). These results imply that the preferential targets of mRNA uridylation vary during seed development or germination. Interestingly, the proportion of mRNAs with a short CDS and a low number of introns is higher among mRNAs that are specifically expressed at the later stages of seed maturation (Supplementary Figure S2C, based on (Le et al. 2010)). These results indicate that mRNAs that accumulate during the late stages of seed maturation are highly uridylated.

**Figure 2:**
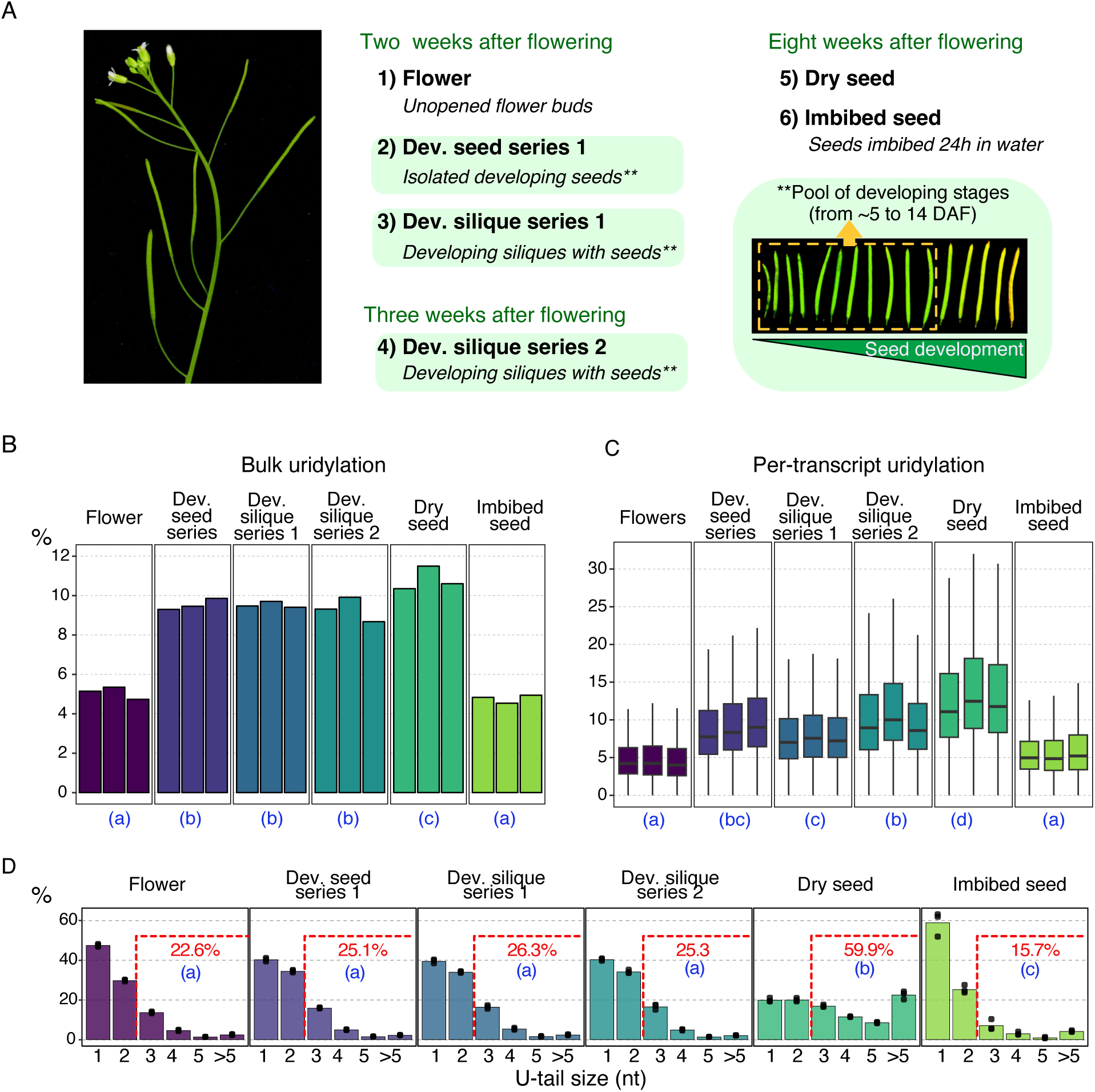
Dynamic of uridylation levels and profiles in seeds. A. Developmental stages used for the preparation of FLEP-seq2 libraries. All samples were harvested for three biological replicates from a minimum of five plants each. B-C. Bulk (B) and per-transcript (C) percentages of uridylation. Per-transcript percentages were calculated for mRNAs detected with at least 50 reads for each replicate. D. % of tails according to the number of Us. Individual points and the bar plot show individual replicates and the average, respectively. The average percentage of tails > 2 Us is indicated above each bar plot. For all plots, significantly different values (adjusted p-value < 0.05, n=3) are labelled by different letters. A generalized linear model for proportion with a quasibinomial distribution was used to compared percentages in (B) and (D) and a linear model was used to compare median in (C).

We have previously shown in Arabidopsis that uridylation targets mostly oligoadenylated mRNAs with A-tail shorter than 25As (Sement et al. 2013; Zuber et al. 2016). Therefore, the high uridylation rates observed in developing seeds and in dry seeds could reflect changes in poly(A) tail populations and could result from an increased proportion of A-tail shorter than 25As. To test this possibility, we analyzed poly(A) size distributions for both non-uridylated and uridylated tails from 10 to 200 As (Figure 3). We also compared the proportion of mRNAs with no or very short tail (tail shorter than 10 As, hereafter called A_<10_-tailed mRNAs) to the proportion of mRNAs with tails between 10 and 25 As (called hereafter A_10-25_-tailed mRNAs) or longer than 25 As (Figure 3). In their study, Jia et al. (Jia et al. 2022) reported tissue-specific regulation of poly(A) tail length with patterns in dry seeds and pollen that differ from other tissues. In line with their results, our analysis confirmed the singularity of the poly(A) profile in dry seeds: the two main peaks of the poly(A) distribution in dry seeds were shifted toward longer tails, from ∼22 and ∼41-46 As in flowers or germinating seeds, to ∼29 and ∼58 As in dry seeds (Figure 3A). Yet, the most striking difference was observed in developing seeds and siliques for which there was a massive shift of poly(A) tail size profiles toward shorter tails compared to other stages (Figure 3A). Our results revealed a strong accumulation of mRNAs with A-tails shorter than 25 As, a small size shift of the first peak to ∼20 nt and a disappearance of the second one. This poly(A) tail size variation was not specific to a subset of mRNAs but instead affected most mRNAs (Figure 4). Altogether, our results reflected important modulations of poly(A) tail sizes during seed development and germination: for the majority of mRNAs, the proportion of A_10-25_-tailed mRNAs was very high in developing seeds, dropped during seed maturation and then increased again at an intermediary level during seed germination (Figure 3 and 4). Actually, the increased proportion of A_10-25_-tailed mRNAs in developing seeds also explained the increased uridylation rate observed at this stage. Indeed, uridylation rates were similar between developing seeds and flowers, when only these A_10-25_-tailed poly(A) tails were taken into account for the calculation of percentages (Figure 3B). Importantly, uridylation rates were higher in dry seeds when compared to other tissues regardless of the poly(A) tail population (Figure 3B) and both short and long uridylated poly(A) tails accumulated (Figure 3A). Altogether, our results revealed two molecular scenarios leading to the accumulation of U-tails in developing and dry seeds. In developing seeds, the higher uridylation rates of mRNAs is likely governed by a stage-specific regulation of poly(A) size distributions. In dry seeds, both the increased uridylation rate and the lengthening of U-tails could reflect changes in the degradation processes during the late stages of maturation such as increased activity of TUTases and/or a slowdown in the degradation of uridylated mRNAs.

**Figure 3:**
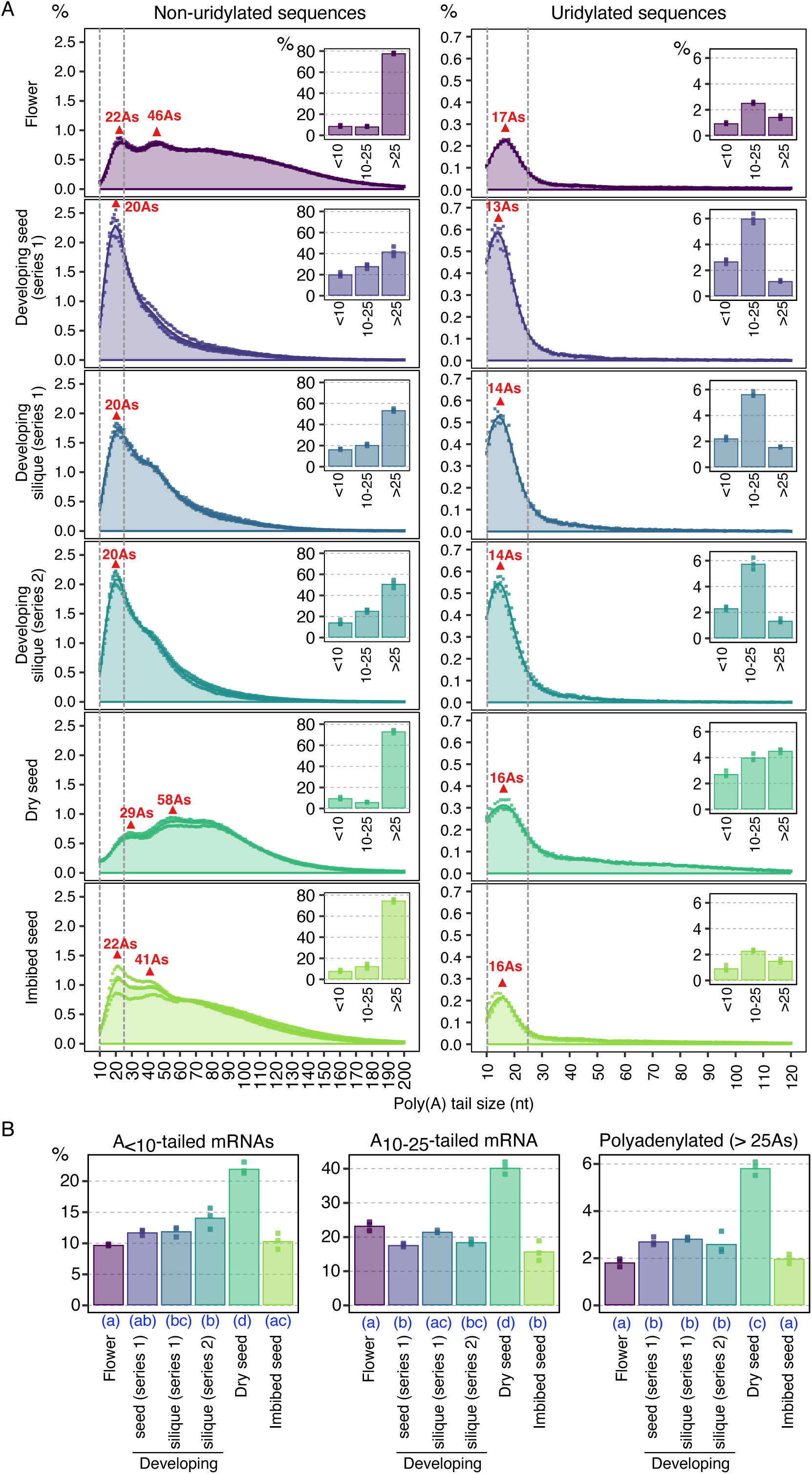
Modulation of poly(A) tail size in seeds. FLEP-seq2 data from developing seeds and siliques (containing seeds), dry seeds and 24h-imbibed seeds. Data are shown for three replicates. A, Distribution of poly(A) tail sizes from 10 to 200 As for non-uridylated (left panel) and uridylated (right panel) reads. Percentages were calculated using the total number of sequences as denominator, including those with A-tails < 10 As. Bar plots next to each distribution show the percentage of reads with no or short A-tail (<10As), with A-tail between 10 and 25 As, or with poly(A) tail longer than 25 As. B, Percentage of uridylation according to A-tail sizes. Percentages were calculated using the total number of sequences of each A-tail population. For all plots of the figure, individual points show the three replicates while color-coded area and bar plots show the average of all replicates. Significantly different values are labelled by different letters (adjusted p-value < 0.05, generalized linear model for proportion with a quasibinomial distribution, n=3).

**Figure 4:**
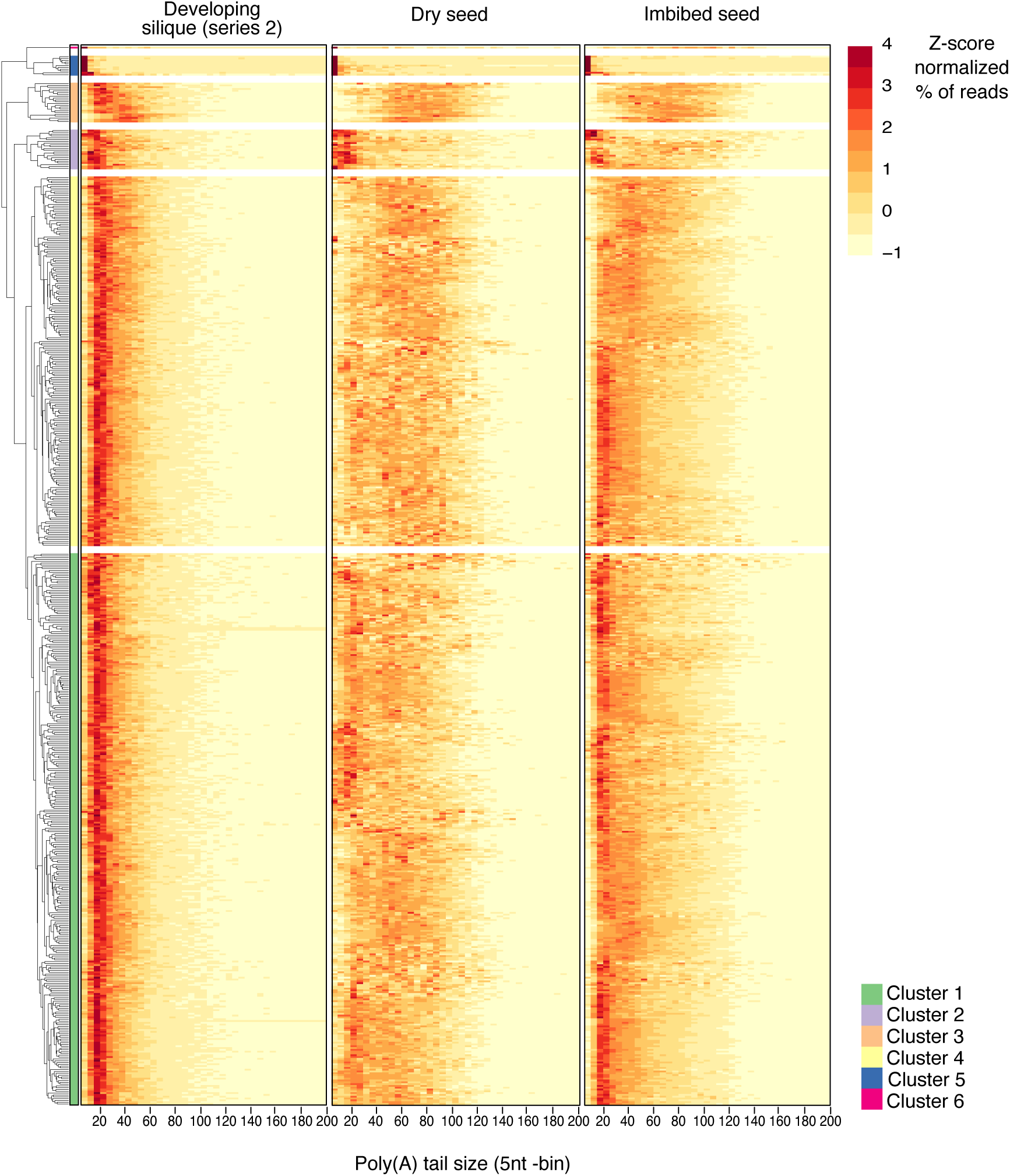
Changes of poly(A) tail size profiles during seed development and germination. Heatmap showing the modulations of poly(A) tail sizes during seed development and germination. % of reads according to poly(A) tail sizes (5nt-bin) were calculated using the total number of sequences as denominator, including those with A-tails < 10 As, and were z-score rescaled. Transcripts were clustered based on their similarity of polyadenylation profiles (k=6).

### Respective contributions of URT1 and HESO1 to mRNA uridylation

While we have previously shown that URT1 is the main TUTase responsible for mRNA uridylation in Arabidopsis leaves and flowers (Sement et al. 2013; Zuber et al. 2016), the contribution to mRNA uridylation of HESO1, the second TUTase characterized in Arabidopsis, has not been studied so far. To apprehend the respective contributions of each TUTase to mRNA uridylation, we compared FLEP-seq2 data from three biological replicates for WT plants (same libraries as described above), *urt1-1*, *heso1-4* and *heso1-4 urt1-1* mutants and for four different tissues: developing siliques, dry mature seeds, germinating seeds and flowers. In total, 48 libraries were analyzed. For each tissue and genotype, we calculated the global uridylation percentage and the uridylation percentages per mRNA if detected with at least 50 reads in all three replicates and four genotypes (Figure 5A, Supplementary Data Set S2). Our results confirmed the preponderant role of URT1 in mRNA uridylation: for all tissues, both global and per-transcript percentages of uridylation strongly decreased in *urt1-1* when compared to WT.

**Figure 5:**
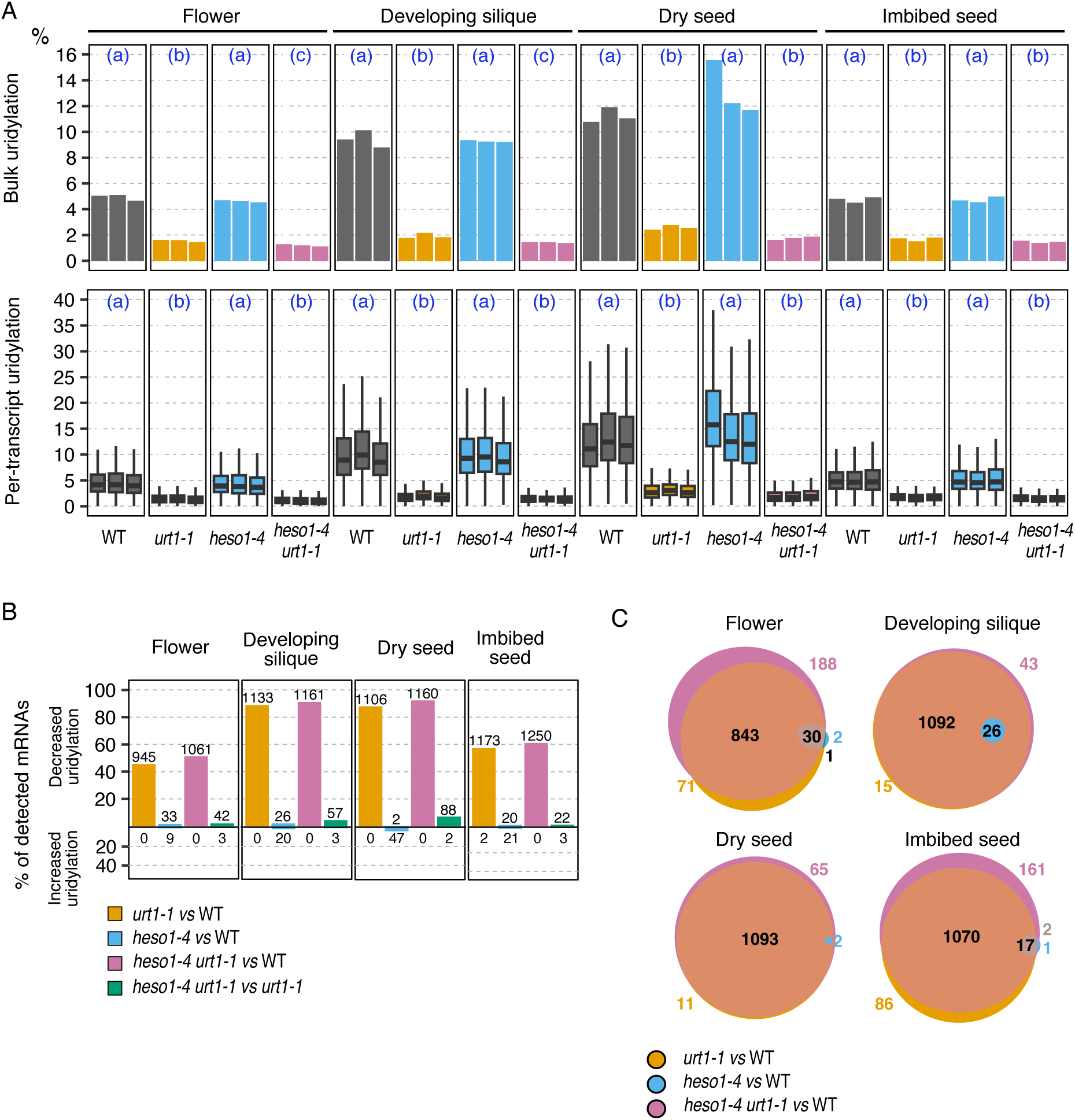
Contribution of URT1 and HESO1 to mRNA uridylation in Arabidopsis. FLEP-seq2 data from developing seeds and siliques (containing seeds), dry seeds and 24h-imbibed seeds for WT, *urt1-1*, *heso1-4* and *heso1-4 urt1-1*. Data are shown for three replicates. Bulk (top panel) and per-transcript (lower panel) percentages of uridylation. Per-transcript percentages were calculated for mRNAs detected with at least 50 reads for each replicate and genotype. Each boxplot displays the median, first and third quartiles (lower and upper hinges), the largest value within 1.5 times the interquartile range above the upper hinge (upper whisker) and the smallest value within 1.5 times the interquartile range below the lower hinge (lower whiskers). Significantly different values are labelled by different letters (adjusted p-value < 0.05, n=3, generalized linear model for proportion with a quasibinomial distribution for the top panel, linear model for the lower panel). B, Proportion of mRNAs differentially uridylated between mutants and WT, or between the *heso1-4 urt1-1* and *urt1-1*. mRNAs that are significantly differentially uridylated were identified using a generalized linear model for proportion with a quasibinomial distribution (adjusted p-value < 0.05, n=3). mRNAs with a significant increase and decrease percentage are shown in the top and lower panel, respectively. Number of mRNAs are indicated above the bars. C, Venn diagrams showing the intersection between the different lists of mRNAs showing a decreased uridylation percentage.

HESO1 contributed only modestly to mRNA uridylation because percentages of uridylation were similar in WT and *heso1-4*, and only further decreased in *heso1-4 urt1-1* when compared to *urt1* by an additional 3.9-8.4% (Supplementary Figure S3A). To identify potential mRNAs preferentially uridylated by HESO1, we performed a statistical analysis and compared mRNAs differentially uridylated between WT and *urt1-1* and *heso1-4* mutants and between *urt1-1* and *heso1-4 urt1-1* mutants (Figure 5B and C, Supplementary Data Set S2). As expected, the highest number of mRNAs with a significant decrease in uridylation rate was found in *urt1-1* and *heso1-4 urt1-1* (Figure 5B) and 92 to 99% of mRNAs with a significant decrease in *urt1-1* were also differentially uridylated in *heso1-4 urt1-1* (Figure 5C). We identified only a few mRNAs showing a significant decrease in *heso1-4* when compared to WT, from 2 in dry seeds to 33 in flowers, or in *heso1-4 urt1-1* when compared to *urt1-1*, from 22 in imbibed seeds to 88 in dry seeds. In addition, we detected a few mRNAs with an increased uridylation rate in *heso1-4* when compared to WT. For example, in dry seeds, 47 mRNAs showed a significant increase in uridylation (Figure 5B). Yet, there was almost no overlap of these gene lists across the different tissues (Supplementary Figure S3B) and no gene ontology or genomic feature was enriched in mRNAs differentially uridylated in *heso1-4* (Supplementary Figure S3C). While URT1 and HESO1 were previously shown to sequentially uridylate some microRNAs (Tu et al. 2015), our analysis did not show any changes in the length of mRNA U-tails in *heso1* mutant and did not demonstrate a similar sequential uridylation for mRNAs (Supplementary Figure S3D). In *heso1-4 urt1-1*, the uridylation percentage median ranged from 0.96 in flowers to 1.7% in dry seeds when considering all reads. When considering only polyadenylated mRNAs (≥ 10 As), for a better comparison with the synthetic DNA fragments previously analyzed, the uridylation percentage median ranged from 0.7% in flowers to 1.1% in imbibed seeds, (Supplementary Data Set S2). These uridylation levels were in the same range as the uridylation background measured for the non-uridylated synthetic DNA fragment (0.6%) (Supplementary Data Set S1). These results suggest that most of the residual uridylation observed in the *heso1-4 urt1-1* likely corresponds to technical background. Altogether, our results showed that URT1 is the main enzyme responsible for the bulk mRNA uridylation regardless of the tissue. HESO1 can participate in mRNA uridylation but that its overall contribution remains modest and limited to a small subset of mRNAs. The existence of a third unidentified TUTase uridylating mRNAs remains elusive at this stage.

### URT1 has a major impact on dry seed transcriptome

In normal growth conditions, no developmental phenotype was observed for *urt1-1*, *heso1-4* or *heso1-4 urt1-1* mutation (Scheer et al. 2021, Supplementary Figure S4A). Yet, the accumulation and the lengthening of mRNA U-tails in dry seeds suggests that uridylation could play an important role in the regulation of gene expression during seed maturation. To evaluate the contribution of both TUTases on the mRNA steady state level in seeds, we produced total RNA-seq libraries from dry seeds of WT, *urt1-1*, *heso1-4* and *heso1-4 urt1-1* plants. For comparison, we produced in parallel RNA-seq from flowers. We performed a differential gene expression analysis to identify mRNAs that are differentially accumulated in mutant seeds and flowers when compared to WT (Figure 6, Supplementary Data Set S3 and S4). Interestingly, the stronger impact of URT1 inactivation on the transcriptome was observed in dry seeds, with 298 and 328 mRNAs differentially accumulated in *urt1-1* and *heso1-4 urt1-1*, respectively (Figure 6A, Supplementary Figure S4B). In flowers, only 32 and 56 mRNAs were differentially accumulated in *urt1-1* and *heso1-4 urt1-1*. These results are in line with the strong uridylation rate observed in seeds and highlight the contribution of URT1-dependent uridylation in shaping the transcriptome of dry seeds, likely during the late stages of seed maturation. Unlike URT1, HESO1 had a stronger impact in flowers with 118 mRNAs differentially accumulated against only 7 in dry seeds. This weaker impact of HESO1 on dry seed transcriptome can be possibly explained by the different expression profiles of the two TUTases during seed development. Indeed, whereas *URT1* is strongly expressed in seeds throughout their development with an increased mRNA accumulation at late maturation stages, the expression of *HESO1* strongly decreased at the end of seed development (Supplementary Figure S4C).

**Figure 6:**
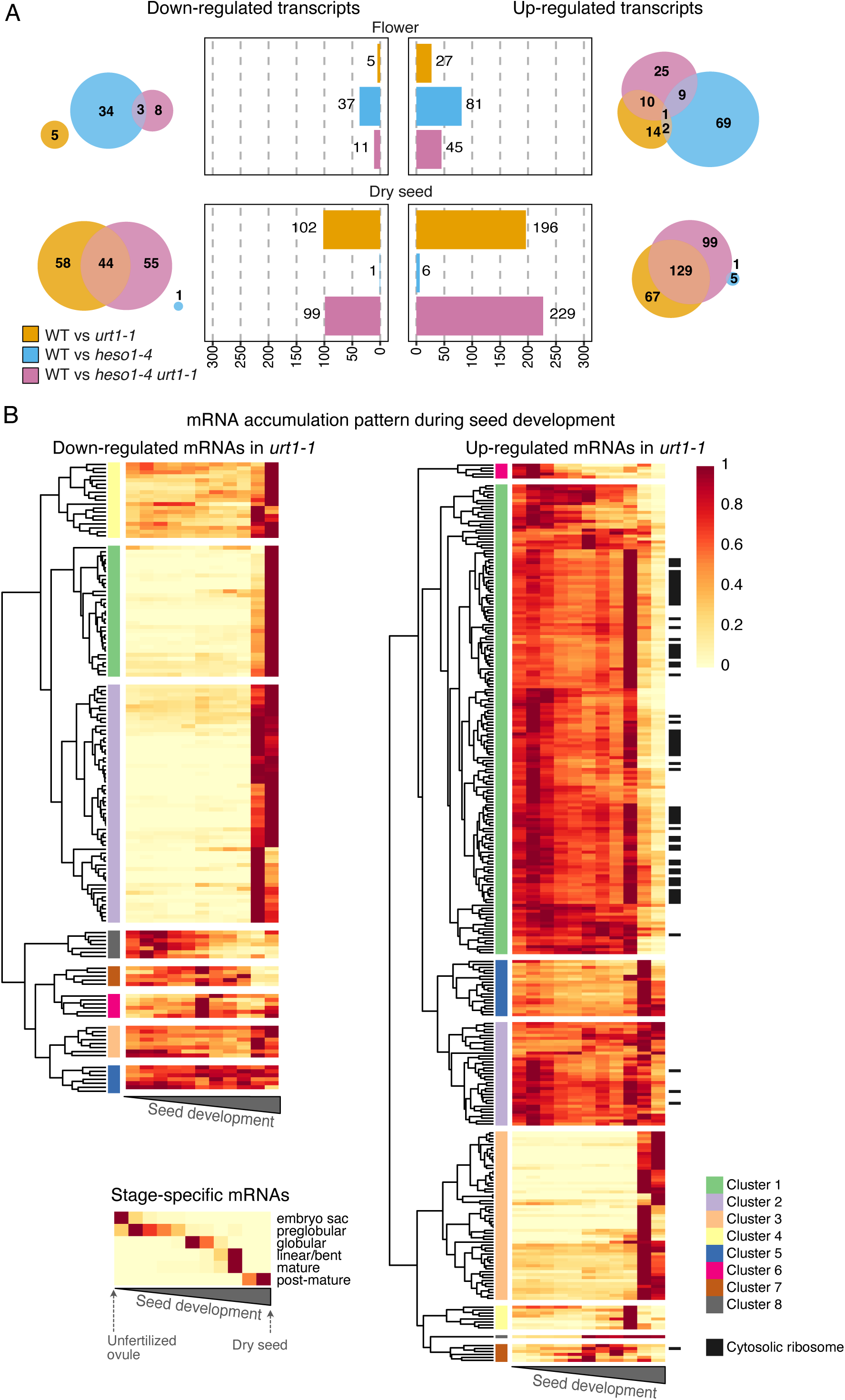
Impact of URT1 and HESO1 inactivation on flower and seed transcriptome. A, Differential expression analysis in flowers and dry seeds of WT, *urt1-1*, *heso1-4* and *heso1-4 urt1-1*. RNAseq data were produced from three biological replicates. Bar plots show the number of significantly differentially regulated transcripts (adjusted p-value < 0.05, DEseq2, n=3) and venn diagrams show the intersection between the different lists. B, Heatmap showing the expression pattern, in a WT context, of transcripts down and up-regulated by URT1 inactivation during seed development, from unfertilized ovules to dry seeds. Expression data are publicly available RNAseq data from Klepikova et al., 2016. As a reference of the different seed development stages, the small heatmap shows the expression pattern during seed development of stage-specific mRNAs (see description in Supplemental Dataset S5). For each mRNA, normalized read count (TMM, EdgeR) were divided by the maximum value of expression level among development stages, so all values vary from 0 to 1. Transcripts were clustered based on their similarity of expression profiles (k=8). Transcripts associated to the cytoplasmic ribosome gene ontology are highlighted by black rectangles.

To search for potential cellular or metabolic processes associated with mRNA uridylation in seeds, we performed gene ontology analysis for the different lists of differentially accumulated mRNAs. Our analysis revealed a strong overrepresentation of gene ontologies associated with ribosome and translation among mRNAs upregulated in both *urt1* and *heso1 urt1* dry seeds (Supplementary Data Set S3). Using publicly available RNA-seq data from Col-0 plants (Klepikova et al. 2016), we then analyzed the expression profile during WT seed development of mRNAs found to be down or up -regulated in *urt1-1* and *heso1-4 urt1-1* seeds (Figure 6B, Supplementary Data Set S5). Interestingly, we observed an opposite trend of expression between mRNAs found to be up and down regulated by URT1 inactivation: while the majority of down-regulated mRNAs (112/145) strongly accumulate at late developmental stages (mRNAs from clusters 1, 2 and 4; Figure 6B left panel), the majority of mRNAs (178/290) up-regulated in *urt1-1* are weakly expressed at the same stages in WT (mRNAs from clusters 1, 4, 6-7; Figure 6B right panel). Of note, most of the up-regulated mRNAs associated with cytosolic ribosomes (64/67) are weakly expressed at the end of seed maturation in a WT context. We then defined two mRNA categories based on their expression patterns during WT seed development: (1) maturation-related mRNAs, which strongly accumulate at the end of seed maturation, and (2) translation-related mRNAs, which encode proteins involved in translation and whose levels decline during late seed maturation (Supplementary Data Set S5). Consistent with previous observations, comparison of these mRNA categories with WT and *urt1-1* RNA-seq data showed that, among the significantly affected mRNAs, maturation-related mRNAs were predominantly downregulated, whereas translation-related mRNAs were exclusively upregulated (Supplemental Figure S4). Altogether, these results suggest that URT1-mediated uridylation promotes the accumulation of maturation-related mRNAs at the end of seed maturation, while participating in the reduction of translation-related mRNAs.

### URT1 inactivation triggers the deadenylation of mRNAs that accumulate during seed maturation

We have previously shown using nanopore Direct RNA sequencing that URT1 shapes poly(A) tail size distributions with a clear shift in the distribution toward short oligo(A) tails in *urt1*-*1* flowers. Our data showed that URT1-dependent uridylation of mRNAs prevents the accumulation of excessively deadenylated mRNAs, likely by both favoring their 5’ to 3’ degradation and hindering deadenylation (Scheer et al. 2021). Our results here confirmed the previous findings: in flowers of *urt1-1*, we observed a clear shift in the global poly(A) tail size distribution compared to WT with an increased accumulation of A_10-25_-tails with sizes centered around 22 As (Figure 7A). While this accumulation of A_10-25_-tails was also clearly visible in imbibed seeds and developing seeds, no change was observed in dry seeds. Yet in all tissues, we observed an increased accumulation of mRNAs with tails shorter than 10 As in *urt1-1* mutant. In flowers, the proportion of A_<10_-tails was moderately increased from 10% in WT to 13% in *urt1-1*. In dry seeds, the increase was more important when compared to other tissues with a proportion going from 12% in WT to 20% in *urt1-1*. These results again reflect the specificity of uridylation-associated processes during the late stages of seed maturation, in which the *urt1-1* mutation did not result in a global accumulation of the A_10-25_-tailed mRNAs but rather triggered an accumulation of excessively deadenylated mRNAs. These results also imply that the function of URT1-mediated uridylation in preventing the accumulation of excessively deadenylated mRNAs could be particularly critical in seeds. Of note, the *heso1-4* mutation did not have a robust impact on the poly(A) tail profiles, regardless of the tissue, and the *heso1-4 urt1-1* double mutation did not worsen the *urt1-1* effect. These results show that HESO1-mediated uridylation of mRNAs is not able to compensate for the absence of URT1 to modulate poly(A) tail size and that both TUTases have distinct functions as previously reported (Tu et al. 2015; Wang et al. 2015; Zuber et al. 2016).

**Figure 7:**
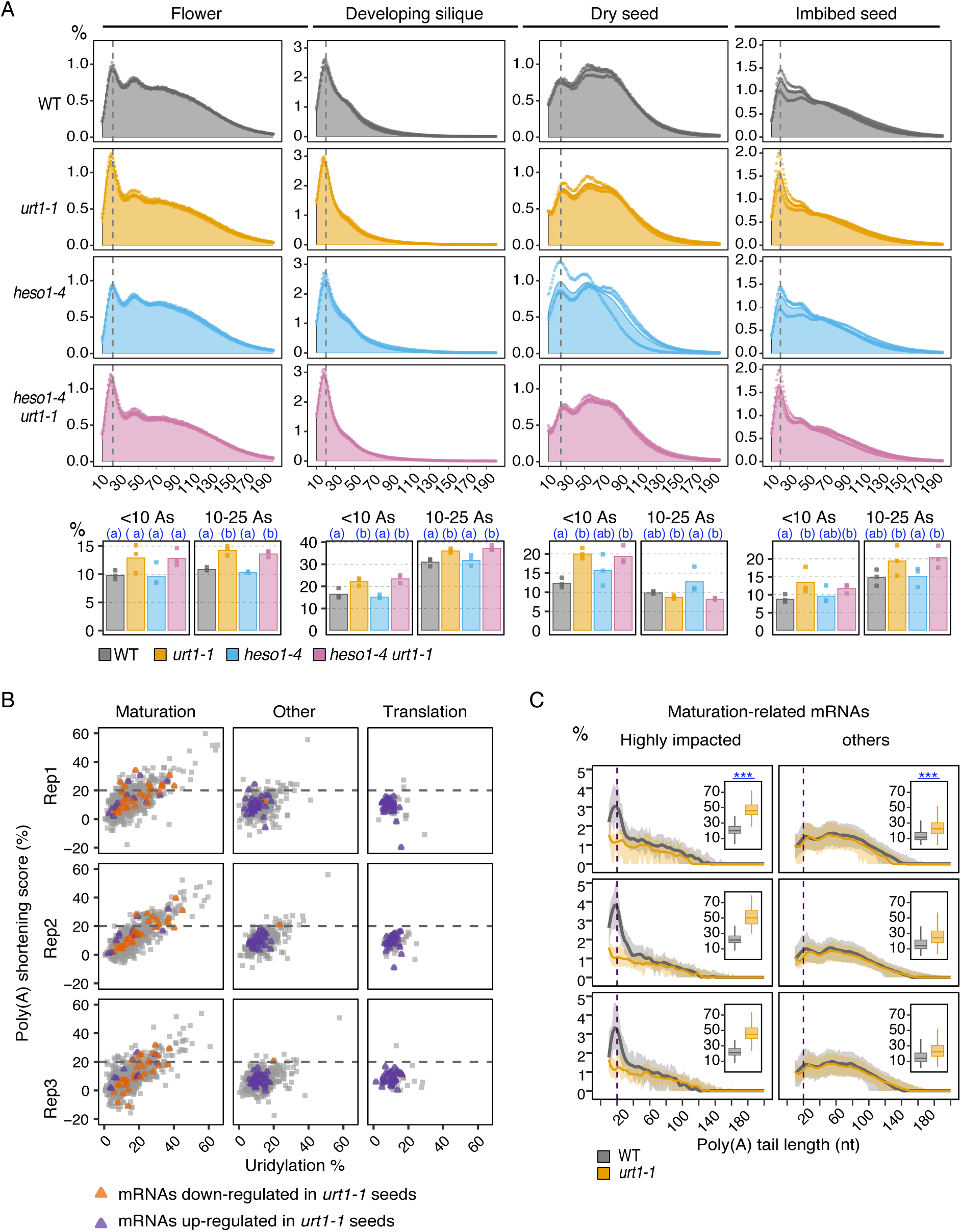
URT1-mediated uridylation prevents a massive accumulation of short A-tails in seeds. FLEP-seq2 data from developing seeds and siliques (containing seeds), dry seeds and 24h-imbibed seeds. Data are shown for three replicates. A, Distribution of poly(A) tail sizes from 10 to 200 As in WT, *urt1-1*, *heso1-4* and *heso1-4 urt1-1*. Percentages were calculated using the total number of sequences as denominator, including those with A-tails < 10 As. Bar plots bellow each distribution show the percentage of reads with no or short A-tail (<10As) or with oligoadenylated A-tail between 10 and 25 As. Individual points show the three replicates while color-coded area or bar plot show the average of all replicates. Significantly different values are labelled by different letters (adjusted p-value < 0.05, generalized linear model for proportion, quasibinomial distribution, n=3). B, Poly(A) shortening scores according to uridylation percentages for maturation- and translation-related mRNAs. Poly(A) shortening score corresponds to the difference in the percentage of A_<10_-tails between *urt1-1* and WT dry seeds. Colored points show mRNAs found to be significantly down-regulated (orange) or up-regulated (purple) in dry seeds by URT1 inactivation. C, Comparison of the poly(A) tail sizes between WT and *urt1-1* for maturation-related mRNAs. Distribution of poly(A) tail sizes from 10 to 200 As. Percentages were calculated using the total number of sequences as denominator, including those with A-tails < 10 As. The lines and the grey shadow show the median and the first and third quartiles of percentages across mRNAs, respectively. Boxplots next to each distribution show the per-transcript proportion of reads with no or short A-tail (<10As). Boxplot displays the median, first and third quartiles (lower and upper hinges), the largest value within 1.5 times the interquartile range above the upper hinge (upper whisker) and the smallest value within 1.5 times the interquartile range below the lower hinge (lower whiskers). Significantly different values (adjusted p-value < 0.05, linear model, n=3) are labelled by stars (* <0.05, *<0.01, ***<0.001).

To identify the most impacted mRNAs in terms of poly(A) shortening in dry seeds, we computed, for each mRNA detected with at least 50 reads, a poly(A) shortening score that corresponds to the difference of the percentage of A_<10_-tails between *urt1-1* and WT dry seeds (Supplementary Data Set S2). These poly(A) shortening scores showed a good correlation with mRNA uridylation: the most uridylated mRNAs also showed the highest scores (Supplementary Figure 5A). Using this poly(A) shortening score, we identified the 93 most impacted mRNAs with an average shortening score exceeding 20%. Interestingly, these 93 mRNAs were significantly enriched in mRNAs encoding proteins related to cellular response to hypoxia and abscisic acid signaling pathway (Supplementary Data Set S2) and most of them (89/93) were genes strongly up-regulated at the end of seed maturation (Supplementary Figure 5B). Moreover, the impact was markedly greater on maturation-related mRNAs compared with translation-related mRNAs (Figure 7B). The maturation-related mRNAs exhibiting a high poly(A) shortening score were characterized by high uridylation rates and shorter poly(A) tails in WT, with a predominant peak in their distribution profile centered at 20 As (Figure 7C). For these highly impacted maturation-related mRNAs, the 20A-peak of the poly(A) distribution profile disappeared in *urt1-1* seeds and the proportion of A_<10_-tails went on average from 21 to 47% (Figure 7C). Among highly impacted maturation-related mRNAs, eight also showed a significant decrease in total mRNA accumulation in *urt1-1* (Figure 7B). This overlap is relatively weak but it shows that for some mRNAs, the poly(A) shortening is coupled with a decrease steady state level of total mRNA. To investigate more carefully a link between the poly(A) shortening and the mRNA steady-state level, we compared per-transcript poly(A) tail profiles for mRNAs down or up-regulated in *urt1-1* or *heso1-4 urt1-1* seeds (Supplemental Figure S5). Our results showed that mRNAs, which show a significant decrease in steady-state mRNA level, were also the most impacted in terms of poly(A) tail size compared to up-regulated or other mRNAs: for mRNAs down-regulated in *urt1-1* dry seeds, the 20 As centered peak of the distribution profile was absent in *urt1-1* dry seeds (Supplemental Figure S5) and the median proportion of A_<10_-tails went from 16-20% in WT to 32-39% in *urt1-1* dry seeds (Supplemental Figure S5). These results show that the poly(A) shortening caused by URT1 defect can trigger a decrease in the mRNA steady-state level at least for some mRNAs.

Altogether our results reveal the diverse consequences of mRNA uridylation during seed maturation: while URT1-mediated uridylation may promote the deregulation of mRNAs encoding translation-related proteins at the end of seed maturation, it may hinder the deadenylation of mRNAs associated with the seed maturation program, thereby facilitating their accumulation.

### mRNAs are subject to differential decay mechanisms during seed maturation

What could explain this antagonistic effect of URT1 inactivation for different population of mRNAs? One explanation could be that translation-related and maturation-related mRNAs experience distinct degradation processes during seed maturation, potentially involving different regulatory factors or subcellular localization. To explore potential variations in their degradation patterns, we analyzed the proportion of 5’ or 3’ fragmented mRNAs in WT and in *urt1-1* for translation and maturation-related mRNAs. FLEP_seq2 allows for the sequencing of full-length RNA molecules. Therefore, a read that does not overlap the first exon or the last exon was considered as 5’- or 3’- fragmented, respectively. When comparing WT and *urt1-1,* we did not observe any significant differences for these two populations of mRNAs (Figure 8A, left panel). Of note, the accumulation of full-length capped transcripts - expected in *urt1-1* due to URT1’s function in promoting decapping - cannot be detected because of current technical limitations of the FLEP-seq2 protocol. Therefore, highlighting an effect of *urt1-1* mutation may require the analysis of a decay mutant (i.e. *xrn4*) where 5’ fragments accumulated. Yet, the most striking observation was the distinct degradation patterns observed in WT between translation and maturation-related mRNAs. Indeed, maturation-related mRNAs were enriched in highly fragmented mRNAs in WT (Figure 8A, right panel) suggesting that these mRNAs are highly decayed during seed maturation. By contrast, translation-related mRNAs in *urt1-1* seeds accumulated only a low proportion of these fragmented forms. These contrasting observations suggest that these two populations of mRNAs exhibit differential degradation efficiency and/or are differentially localized. Interestingly, maturation-related mRNAs were also characterized by: higher uridylation rates (Figure 8B), longer U-tails (Figure 8C) and higher accumulation of very short A-tails (Figure 8D) in WT seeds when compared to translation-related mRNAs. These three characteristics resemble those previously observed in a *xrn4* mutant (Sement et al. 2013; Nagarajan et al. 2019), suggesting that maturation-related mRNAs are less efficiently targeted by the 5’-3’ degradation pathway. Altogether, these results suggest that mRNA associated to translation and those involved in the seed maturation program are subject to distinct decay processes and are differentially accessible to the 5’-3’ degradation pathway during the late stages of seed maturation.

**Figure 8.**
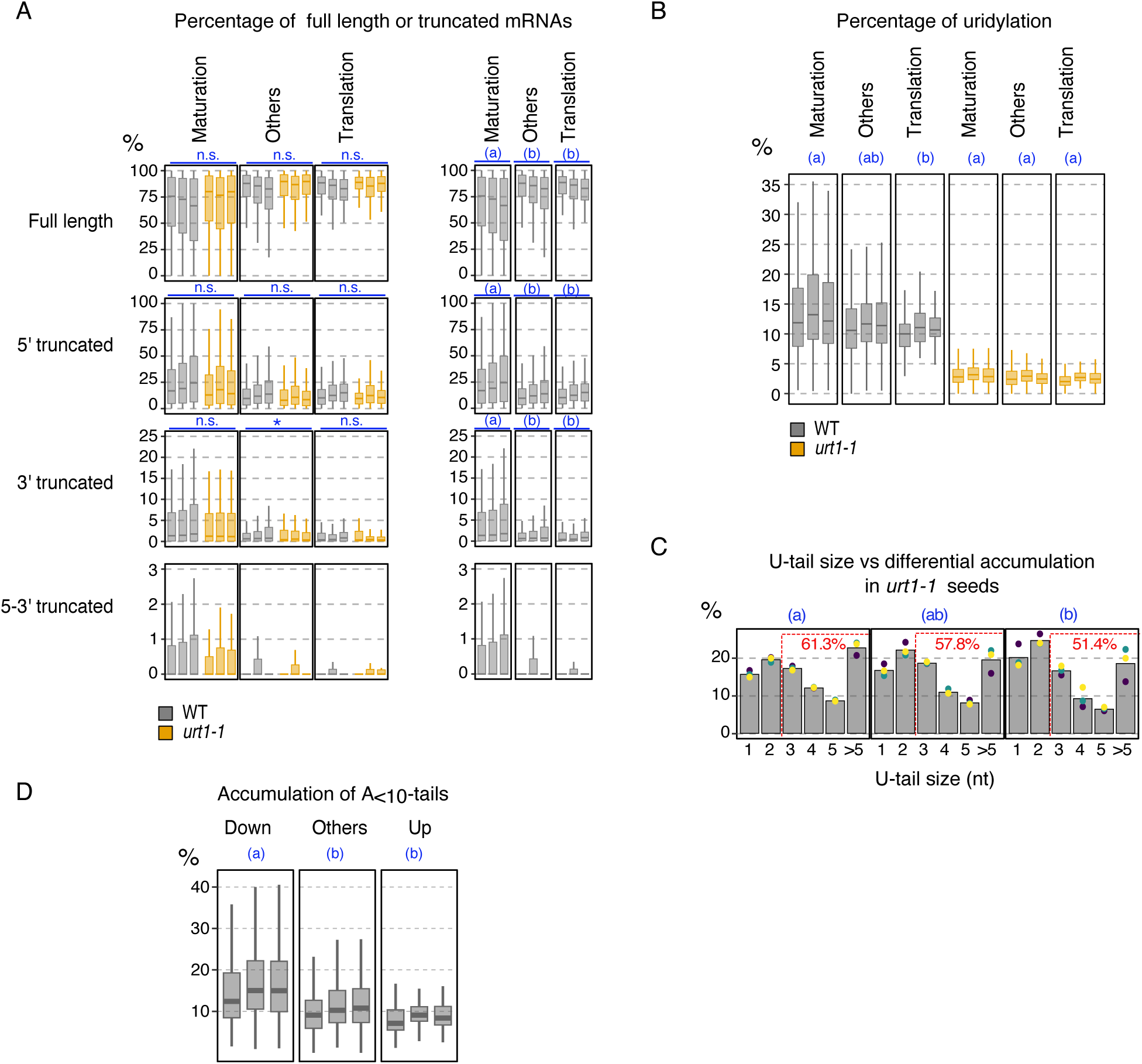
mRNAs are subject to differential decay processes in seeds. A, Percentage of full-length and truncated mRNAs for maturation- and translation-related mRNAs. On the left panel, percentages are shown for all tails in WT and in *urt1-1*. On the right panel, results are shown only for all tails in WT to allow direct comparison between maturation-and translation-related mRNAs. Significantly different values (adjusted p-value < 0.05) are labelled by stars (* <0.05, *<0.01, ***<0.001) or by different letters for multiple comparison (linear model, n=3). B, Per-transcript percentage of uridylation for maturation- and translation-related mRNAs. Significantly different values (adjusted p-value < 0.05) are labelled by different letters (linear model, n=3). C, % of tails according to the number of Us for maturation and translation-related mRNAs. Individual points and the bar plot show individual replicates and the average, respectively. The average percentage of tails > 2 Us is indicated above each bar plot. Significantly different percentages (adjusted p-value < 0.05) are labelled by different letters (generalized linear model for proportion, quasibinomial distribution, n=3). D, Per-transcript percentages of A_<10_-tails in WT. Significantly different values (adjusted p-value < 0.05) are labelled by different letters (linear model, n=3). In (A), (B), (D), each boxplot displays the median, first and third quartiles (lower and upper hinges), the largest value within 1.5 times the interquartile range above the upper hinge (upper whisker) and the smallest value within 1.5 times the interquartile range below the lower hinge (lower whiskers).

### URT1 inactivation alters primary dormancy levels of seeds

The important contribution of URT1-dependent uridylation in shaping the seed transcriptome prompted us to investigate a possible impact of URT1 inactivation on seed germination. We first monitored the germination kinetic of after-ripened seeds, in which primary dormancy is alleviated. We also applied different duration of accelerated aging treatment to investigate a potential impact of URT1 inactivation on seed longevity. Our results did not reveal any impact of URT1 inactivation on the germination kinetic of after-ripened seeds and the final germination rates or the germination speed remained unchanged in *urt1-1* mutant even after accelerated aging treatments (Supplementary Figure S6A). We then conducted germination assays on freshly harvested seeds, which usually show high levels of dormancy, and we monitored the percentages of germination after 7 days. These assays revealed a significant decrease in primary dormancy in *urt1-1* mutant when compared to WT (Figure 9A). This weaker dormancy of *urt1-1* freshly harvested seeds was repeatedly observed for seeds harvested from individual plants and from individual batches of plants (at least four plants for four individual batches, Figure 9A). We performed additional assays by including *heso1-4* and *heso1-4 urt1-1* mutants, *urt1-2* a second allele for *urt1* mutant (Sement et al. 2013) and as control, *dog1-4* (non-dormant control, knock-out mutant (Cyrek et al. 2016) and *dog1-5* (dormant control, mutant overexpressing *DOG1* (Fedak et al. 2016). While the lower level of primary dormancy was confirmed for *urt1-2* and in *heso1-4 urt1-1* seeds, the level of dormancy was not affected in *heso1-4* (Supplementary Figure S6B and C). These results indicate that the uridylation function in the regulation of primary dormancy is specific of the URT1-mediated uridylation. Interestingly, *DOG1* a regulator of seed dormancy, whose protein level was shown to highly correlate with dormancy (Nakabayashi et al. 2012), was among the most uridylated mRNAs and also among the most impacted mRNAs in terms of poly(A) shortening in *urt1-1* seeds (Figure 9B and C). The proportion of A<10-tailed mRNAs significantly increased from 24% in WT to 47% in *urt1-1* seeds (Figure 9C). In addition, although not significantly changed in after-ripened seeds (Figure 9D), the level of total *DOG1* mRNA was significantly reduced in *urt1-1* freshly harvested seeds (Figure 9E). Therefore, the weaker dormancy levels in *urt1-1* seeds could be a consequence of increased deadenylation and subsequent destabilization of *DOG1* mRNAs. The weaker dormancy could also be explained by the increased accumulation in *urt1-1* seeds of translation-related mRNAs, as an up-regulation of genes encoding protein of the translational machinery was previously linked to a loss of dormancy (Cadman et al. 2006; Nelson and Steber 2017; Buijs et al. 2020). In any case, our results highlighted the importance of URT1-mediating uridylation in the regulation of primary dormancy.

**Figure 9:**
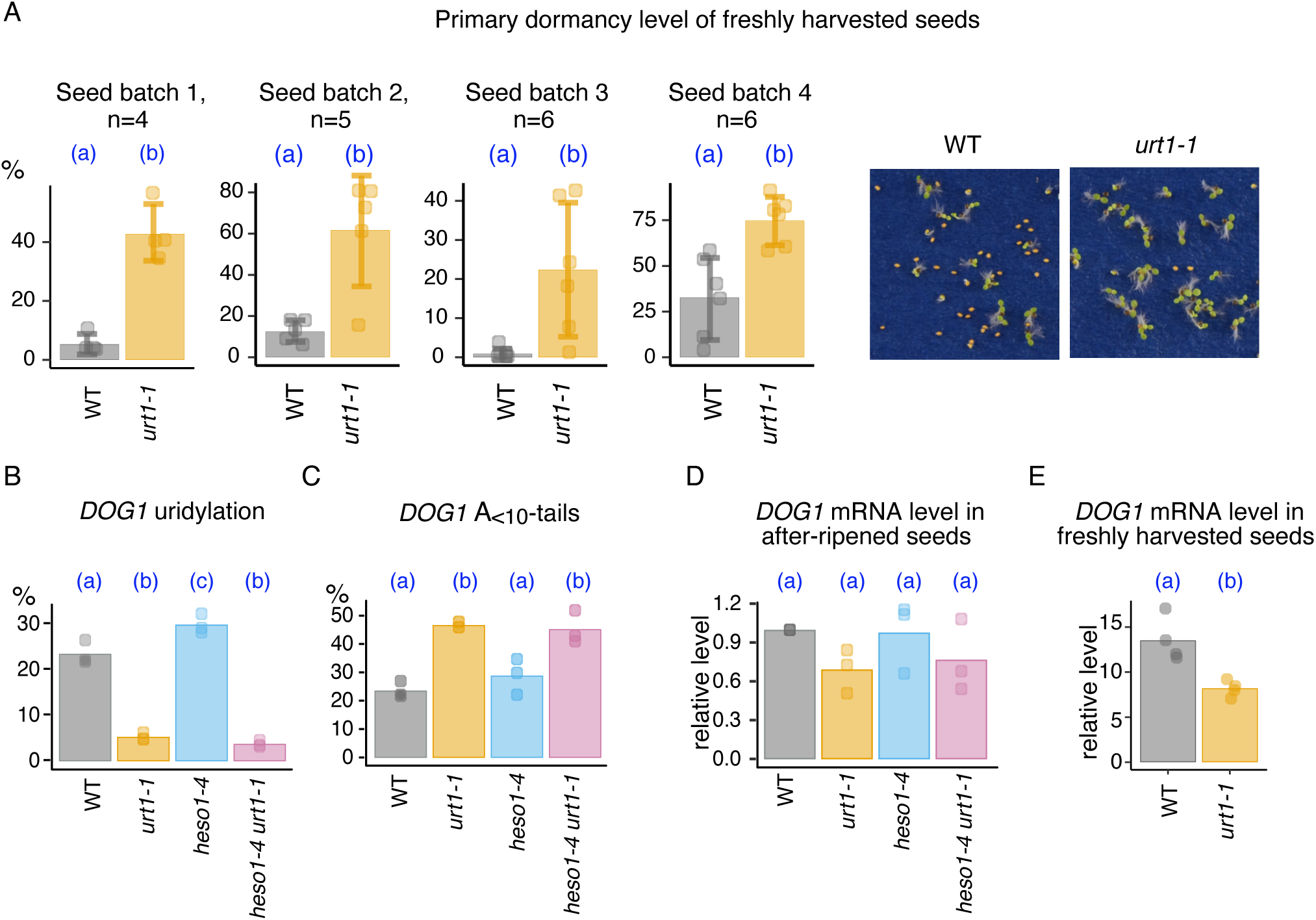
URT1 inactivation alters primary dormancy level. A, Level of primary dormancy of WT and *urt1-1* freshly harvested seeds for four independent batches of seeds. The bar plots show the average percentage of germinated seeds after seven days and individual points show the percentages for seeds harvested from individual plants. Error bars indicate the standard deviation. Significantly different values are labelled by different letters (p-value < 0.05, generalized linear model for proportion, quasibinomial distribution, n indicated on each graph). A representative image of germination assays is shown on the right. B, % of uridylation of the *DOG1* mRNA in dry seeds of WT, *urt1-1*, *heso1-4* and *heso1-4 urt1-1* based on FLEP-seq2 data (n=3). C, % of A_<10_-tailed mRNAs of the *DOG1* mRNA in dry seeds of WT, *urt1-1*, *heso1-4* and *heso1-4 urt1-1* based on FLEP-seq2 data (n=3). In (B) and (C), individual points show biological replicates and bar plots show the average. Significantly different values are labelled by different letters (adjusted p-value < 0.05, generalized linear model for proportion, quasibinomial distribution, n=3). D, Relative expression of *DOG1* (normalized to WT) in after-ripened seeds of WT, *urt1-1*, *heso1-4* and *heso1-4 urt1-1* based on RNAseq data. Individual points show biological replicates and bar plots show the average. Significantly different values are labelled by different letters (adjusted p-value < 0.05, DEseq2, n=3). E, Relative expression of *DOG1* (normalized to the reference gene *UBIQUITIN-CONJUGATING ENZYME 21*) in freshly harvested seeds of WT and *urt1-1*. Individual points show seeds harvested from independent plants and bar plots show the average. Significantly different values are labelled by different letters (p-value < 0.05, two-tailed Wilcoxon rank-sum test for unpaired data, n=4).

**Figure 10:**
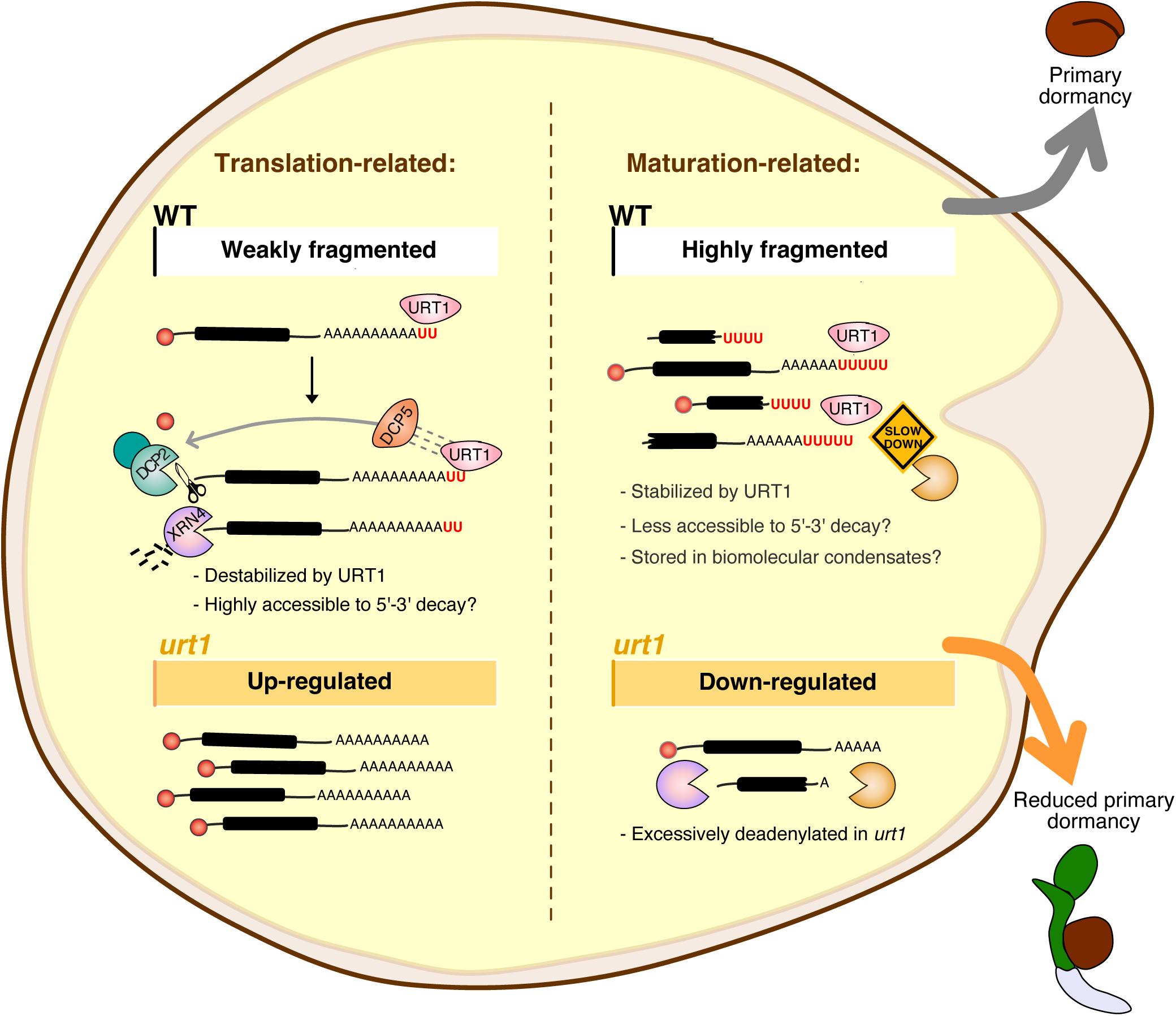
Model of URT1 dual functions during the late stages of seed maturation. We propose that the dual roles of URT1-mediated uridylation, promoting mRNA decay versus stabilizing transcripts, results from the profound changes in RNA metabolism at the end of seed maturation. On one hand, translation-related mRNAs exhibit low levels of fragmentation and may be efficiently degraded by 5′-to-3′ exonucleolytic decay. These transcripts are destabilized by URT1-mediated uridylation and are therefore up-regulated in a *urt1* mutant. On the other hand, maturation-related mRNAs are highly fragmented and may be less accessible to 5′-to-3′ decay. We propose that these mRNAs are stored in biomolecular condensates. These transcripts are stabilized by URT1-mediated uridylation and are consequently down-regulated in a *urt1* mutant.

## Discussion

Developmental transitions that occur in the seed imply extensive changes of gene expression, at both transcript and protein levels, which have been documented thoroughly in Arabidopsis during seed development (Le et al. 2010; Belmonte et al. 2013; Bai et al. 2023) or germination (Nakabayashi et al. 2005; Narsai et al. 2011; Basbouss-Serhal et al. 2015; Bai et al. 2017). While transcriptional regulation has been well studied, the post-transcriptional mechanisms involved during seed development and germination remain less understood, especially those related to RNA degradation. mRNA uridylation is a widespread post-transcriptional modification that influences mRNA decay. In the present work, we optimized the data analysis of the nanopore sequencing-based FLEP-seq2 method developed by (Jia et al. 2022) to generate a first inventory of mRNA uridylation events in different Arabidopsis tissues, including developing and germinating seeds. Our results showed overall high levels of mRNA uridylation in developing and in dry seeds, higher than in all other analyzed tissues. In developing seeds, the high uridylation rates of mRNAs were associated with a strong regulation of poly(A) tail size and likely resulted from the massive accumulation of A_10-25_-tailed mRNAs that are the preferential targets of mRNA uridylation (Sement et al. 2013; Zuber et al. 2016). By contrast in dry seeds, high uridylation rates were independent of a poly(A) tail size regulation and reflected either an increased activity of URT1 and/or a decrease degradation of uridylated mRNA during the late stages of maturation. *URT1* mRNA strongly accumulates at the end of seed maturation suggesting that the uridylation activity could be particularly high at this stage. Yet, the lengthening of U-tails observed specifically in dry seeds could also reflect a lower degradation activity. Indeed, a similar lengthening was previously shown when 5’ to 3’ or 3’ to 5’ degradation was impaired (Sement et al. 2013; Lim et al. 2014), but not when URT1 was overexpressed (Scheer et al. 2021). The maturation of orthodox seeds is characterized by a gradual decrease in water content coupled with a strong reduction of the metabolic activity. Therefore, the accumulation of long U-tails during the late stages of seed maturation could reflect the gradual shutdown of the enzymatic degradation pathways, which would be spatiotemporally uncoupled from uridylation processes for reasons yet to be investigated. While the uridylation level of mRNAs was maximal in dry seeds, it was reduced during germination after 24 hours of imbibition in water. The size of U-tails was also shorter in imbibed seeds. This quantitative and qualitative change in uridylation profiles of mRNAs likely reflects the resumption of degradation processes during early germination stages.

These high mRNA uridylation levels in the seed were coupled with a greater impact of URT1 inactivation on the transcriptome of dry seeds as compared to flowers (the present study) or leaves (Sement et al. 2013). These results revealed the importance of URT1-dependent uridylation in the regulation of gene expression during the late stages of seed maturation, which therefore constitutes a relevant model to dissect the multiple consequences of mRNA uridylation on RNA metabolism. The housekeeping function of mRNA uridylation in Eukaryotes is to facilitate mRNA degradation. In plants, we have previously shown that mRNA uridylation prevents excessive deadenylation thereby favoring the polarity of degradation from 5’ to 3’. This dual function of mRNA uridylation is not limited to plants as a recent study shows that a defect in uridylation also leads to an excessive deadenylation in fission yeast (Grochowski et al. 2024). Our results here highlighted the respective contribution of the dual functions of URT1-dependent uridylation in regulating distinct subsets of the transcriptome at the end of seed maturation. On one side, we showed that URT1-mediated uridylation promotes the deregulation of translation-related mRNAs during seed maturation. Translation is developmentally regulated in maturing seeds: the overall abundance of ribosomes increases during the initial phase of seed maturation, but then decreases during seed desiccation, indicating a general decline in translation (Bai et al. 2023). Interestingly, our results show that translation-related mRNAs exhibited a significantly higher level of uridylation in seeds compared to other tissues showing that these mRNAs are important targets of uridylation at this developmental stage. Therefore URT1-mediated uridylation could participate in the down-regulation of the translation by facilitating the elimination of translation-related mRNAs that become unnecessary when the seed enters a quiescent state. On the other side, we showed that URT1-mediated uridylation facilitates the accumulation of a hundred of mRNAs that typically accumulate at the end of seed maturation, likely by protecting them from an excessive deadenylation. In the absence of URT1, the decrease accumulation of these mRNAs was coupled with an important shortening of their poly(A) tail size: the population of mRNAs with tails centered around 20 As decreased in favor of

a significant accumulation of mRNAs with very short tails. This distribution peak centered around 20 As may reflect the fingerprint of one Poly(A) Binding protein (PABP). Indeed, even if PABPs was shown to occupy ∼ 27 As (Baer and Kornberg 1983), the minimal A-tail size for high affinity binding *in vitro* is 12 As (Sachs et al. 1987). Moreover, Arabidopsis PABPs were shown to bind mRNAs with tails as short than 16 As, as well as to uridylated mRNAs (Zuber et al. 2016). In fact, our previous findings revealed that URT1-mediated uridylation can repair deadenylated ends to restore a binding site for PABP (Zuber et al. 2016), supporting the idea that in seeds URT1-mediated uridylation could facilitate the binding of the PABP to mRNAs associated with the maturation program, thereby stabilizing them.

Dry seeds are characterized by the accumulation of many mRNAs at the end of maturation, called stored mRNAs or long-lived mRNAs (Sajeev et al. 2019; Sano et al. 2020). Some of these stored mRNAs were shown to serve as templates for de novo translation during germination (Rajjou et al. 2004; He et al. 2011; Sano et al. 2020), while the others may be reminiscent of the maturation program, thereby reflecting the transcriptomic regulations taking place on the mother plant. We propose that the antagonistic roles of URT1-mediated uridylation - promoting mRNA decay versus stabilizing transcripts – are explained by the profound changes in RNA metabolism at the end of seed maturation, which lead to the storage of thousands of mRNAs. Interestingly, the analysis of degradation patterns in WT suggests that translation-related and seed maturation-related mRNAs undergo distinct decay process. While translation-related mRNAs may be actively degraded from 5’ to 3’, mRNAs associated with the maturation program may be less accessible to the 5’ to 3’ decay pathway, thereby enabling their storage. The mechanisms underlying the storage of these mRNAs in dry seeds remain largely unknown. Processing bodies and stress granules, biomolecular condensates balancing mRNA degradation, storage and translation (for reviews in plants Kearly et al., 2024; Solis-Miranda et al., 2023), were proposed to serve as a storage location and to act as a protective armor (Sajeev et al. 2019). Interestingly, URT1 is a component of both processing bodies and stress granules (Sement et al. 2013) and possesses an N-terminal intrinsically disordered region (IDR) that is conserved across the plant cell lineage (Scheer et al. 2021). Given the central role of IDRs in condensate formation (Shi et al. 2025), this suggests that URT1 localization to biomolecular condensates may be a conserved feature among plant URT1 orthologs. We therefore hypothesized that seed stored-mRNAs could escape the main 5’ to 3’ pathway by being sequestered in biomolecular condensates. This recruitment to biomolecular condensates could be a way to protect stored mRNAs from degradation similarly to what was shown in response to heat-stress for some heat response-related transcripts that are recruited to stress granules by ALBA proteins (Tong et al. 2022). Plant processing bodies were also proposed to function as mRNA-reservoirs in dark-grown seedling, regulating the selective translation of mRNAs encoding proteins needed for photomorphogenesis (Jang et al. 2019).

URT1-mediated uridylation shapes the transcriptome of dry seeds. But what are the physiological consequences of this URT1 function? The present work identified URT1 as a novel positive regulator of the primary dormancy. The dormancy is an important seed agronomical trait that is defined by the state in which seeds are unable to germinate despite having favorable environmental conditions. While secondary dormancy is induced when imbibed non-dormant seeds are exposed to adverse environmental conditions, primary dormancy is established during seed development (for reviews about dormancy see Chahtane et al., 2017; Née et al., 2017; Sajeev et al., 2024). A previous study already highlighted the importance of RNA degradation pathways in the regulation of primary dormancy. In their work, the authors showed primary dormancy phenotypes for two major components of the 5’ to 3’ decay pathways, the decapping enhancer VARICOSE (VCS) and the main 5’ to 3’ exoribonuclease EXORIBONUCLEASE4 (XRN4), with *vcs* and *xrn4* mutant exhibiting reduced and higher dormancy level, respectively (Basbouss-Serhal et al. 2017). Explaining these opposite phenotypes, the authors identified subsets of mRNAs specifically targeted by VCS or XRN4 that positively regulate the germination or on the contrary are involved in the maintenance of dormancy, respectively (Basbouss-Serhal et al. 2017). The varying influence of these two 5’-3’ decay factors show the complexity of the interplay between RNA degradation and the regulation of the primary dormancy. The regulatory role of URT1 could occur during the late stages of seed maturation, when primary dormancy is established, or during seed imbibition, when dormant seeds arrest the germination program. The high uridylation rate of mRNAs in dry seeds and its drop as soon as 24 hours after imbibition support the first hypothesis. Moreover, the transcriptomic changes in *urt1-1* dry seeds could also explained the reduced depth of primary dormancy triggered by URT1 inactivation. Indeed, our transcriptomic analysis highlighted a reduced accumulation of polyadenylated mRNAs for DOG1 and for genes related to the ABA signaling pathway and an increased expression of translation-related mRNAs in *urt1-1* seeds. Altogether, our results suggest that of URT1-mediated uridylation could positively regulate the establishment of the primary dormancy during seed maturation. The depth of dormancy is controlled by the environmental conditions experienced by the mother plant, for example by the temperature or by the nitrate availability (Klupczyńska and Pawłowski 2021). A better understanding of how plants perceive and signal these environmental factors to modulate the depth of seed dormancy is a major challenge from an agronomic perspective. In the future, it would be particularly interesting to explore the contribution of mRNA uridylation and RNA decay pathways in shaping the seed transcriptome in response to these environmental signals and to investigate its influence on the regulation of dormancy.

## Methods

### Plant material and growth condition

The *Arabidopsis thaliana* plants used in this work are of Columbia accession (Col-0). Arabidopsis mutants analyzed in this study are T-DNA insertion lines described previously: *urt1-1* (SALK_087647C) (Sement et al. 2013), *urt1-2* (WISCDSLOXHS208_08D), *heso1-4* (GK-369H06-017072) (Joly et al. 2023; Vigh et al. 2024), *dog1-4* (Cyrek et al. 2016) and *dog1-5* (Fedak et al. 2016). Arabidopsis plants were grown on soil in long-day conditions with a 16 h light/8 h darkness photoperiod in a growth chamber at 21/18 °C (neon light, 110-115 µsm). The flowers (unopened flower buds), developing siliques (pool of siliques from ∼5 to 14 DAF before visible yellowing is apparent) and dry mature seeds for RNAseq and FLEP-seq2 were harvested from a minimum of five plants each for three biological replicates. Samples were harvested two weeks after the onset of flowering for flowers and the first series of developing siliques, three weeks after the onset of flowering for the second series of siliques, and finally eight weeks after the onset of flowering for dry mature seeds, when all siliques of the plants were dry. For the first series of developing siliques, developing seeds were also isolated from their encapsulating siliques using liquid nitrogen and dry ice as described in (Bates et al. 2013). 24-hour imbibed seeds were obtained after imbibition of the dry mature seeds for 24 hours in milliQ water (16 h-day at 21°C / 8 h-night at 18°C cycles).

### Germination test and accelerating aging treatment for after-ripened seeds

After-ripened seeds were harvested 7 to 8 weeks after the onset of flowering. Accelerating aging treatment of seeds were then performed according to (Rajjou et al. 2008). All seeds were equilibrated for three days at 85% relative humidity (20°C) in a constant climate chamber. Day-0 controls were immediately dried back for three days at 32% relative humidity (20°C) and stored at 4°C. Aging treatment was then done by storing the seeds for two, four or seven days at 85% relative humidity (40°C). After each treatment, seeds were dried back for three days at 32% relative humidity (20°C) and stored at 4°C. The germination tests were then performed by placing plates under constant light (20°C) and germination was scored every day for four days. Source data for germination assays are shown in Supplementary Data Set S6.

### Primary dormancy assays

Freshly harvested seeds used for primary dormancy assays were obtained from plants grown in a greenhouse under a long-day photoperiod at 22°C. Importantly, the dry, mature seeds were collected when flower buds were still on the main stem. Dormancy assays were repeated four times for four independent batches of plants. Seeds were collected from four to six individual plants for each independent experiment and each genotype. Approximately 100 to 200 seeds for each plant were sown on blue germination paper (Hoffman Manufacturing Inc.) supported by a layer of thick fabric saturated with water. Plates were placed in a greenhouse under a long-day photoperiod at 22°C and germination was scored after seven days (Figure 9) or four days (Supplementary Figure S6). Source data for germination assays are shown in Supplementary Data Set S6.

### Oligonucleotides

Oligonucleotides used in this study are listed in Supplementary Data Set S7.

### RNA isolation for FLEP-seq2 and RNA-seq

Total RNA from seeds (dry, imbibed and developing seeds) and siliques was extracted using liquid nitrogen according to the protocol (Meng and Feldman 2010) except for the last precipitation performed at -80 °C for 1 hour. RNA from flowers was extracted using liquid nitrogen with TRI Reagent (EUROMEDEX, TR118-100) following the manufacturer’s protocol. Total RNA was then purified using the RNeasy MinElute Cleanup kit (Qiagen, 74204).

### Library preparation and nanopore sequencing

FLEP-seq2 libraries were prepared from 3 µg of total RNA as described in (Long et al. 2021; Jia et al. 2022). Briefly, 3 µg of RNA were ribodepleted using the Arabidopsis thaliana riboPOOL kit (siTOOLSs BIOTECH, dp-K024-000008) and then ligated to 50 pmol of a 5’-riboadenylated DNA oligonucleotide (Universal miRNA cloning linker, NEB, Supplementary Data Set 7) in a 20 µl reaction containing 20 U of T4 RNA Ligase 2 truncated KQ (NEB, M0373S), 1X T4 RNA Ligase Reaction Buffer (NEB, 50 mM Tris-HCl pH 7.5, 10 mM MgCl2, 1 mM DTT), 25% PEG8000 and 40 U RNAseOUT (Life Technologies, 10777019). Ligation reactions were incubated for 10 hours at 16°C. The product was cleaned up using RNA Clean & Concentrator-5 kit (ZYMO, R1013). The cDNA synthesis was then performed in two steps. First, the ligated RNAs were incubated with 2 pmol of a custom reverse transcriptase primer (FLEP_seq2_RT, Supplementary Data Set 7) and 10 nmol of dNTPs, incubated at 65°C for 5 min, and placed on ice for 2 min. Then, 1X Maxima RT Buffer (Thermo Fisher, EP0752), 40 U RNaseOUT, 20 pmol of Strand-Switching Primer (SSPI or SSPII, ONT, SQK-PCB111.24) and 200 U Maxima H Minus Reverse Transcriptase (Thermo Fisher, EP0752) were added to the reaction in a 20 µl final volume. Samples were incubated in a thermal cycler at 42°C for 90 min 4°C; 10 cycles at 50°C for 2 min and 42°C for 2 min; and a final step at 85°C for 5 min. cDNA libraries were amplified for 14 cycles in a 25 µl reaction containing 1X PrimeSTAR GXL Buffer (TaKaRa, R050A), 5 nmol of each dNTP, 7.5 pmol of each barcoded primer of the PCR-cDNA Barcoding kit (ONT, SQK-PCB111.24) and 0.625 U of PrimeSTAR GXL DNA polymerase (TaKaRa, R050A). The PCR conditions were as follows: one step at 98°C for 30 s; 14 cycles at 98°C for 10 s, 65°C for 15 s and 68°C for 10 min; a final step at 68°C for 5 min. The PCR products were then incubated with 20 U of Exonuclease I (NEB, M0293) at 37°C for 15 min, followed by a step at 80°C for 15 min. For each sample, 4-6 PCR reactions were carried out and subsequently pooled. The PCR products were cleaned up with three successive rounds of purifications with NucleoMag NGS Clean-up and Size Select kit (Macherey-Nagel, 744970.50) using bead ratios of 0.8X, 0.55X and 0.55X, respectively. Libraries were quantified using the Quant-iT HS DNA assay (Invitrogen, Q33120) and QuBit fluorometer, and the size and the quality were checked using the Agilent High Sensitivity DNA Kit and the Agilent 2100 Bioanalyzer System. Finally, after adding nanopore adapter (Rapid Adapter T, SQK-PCB109/SQK-PCB111), libraries were sequenced for ∼72 hours on a MinION or a P2 PromethION 2 Solo device using a FLO-MIN106 (R9.4.1) or a FLO-PRO002 (R9.4.1) flowcell, respectively.

### Computational analysis for FLEP-seq2

Raw signal FAST5 files were basecalled using Guppy (Version 6.3.8+d9e0f64) with default parameters using the high accuracy model (-c dna_r9.4.1_450bps_hac). The reads with a mean quality score inferior to 7 were filtered out. The remaining reads were mapped to the *Arabidopsis thaliana* TAIR10 genome using Minimap2 (2.26-r1175) with the Long-read spliced alignment preset enabled (-ax splice) without keeping secondary alignments (--secondary=no). The detection of the 3’ ends of the transcripts and the determination of their length have been performed using a pipeline adapted from (Jia et al. 2022) and available on Github (https://github.com/jackson-peter/FLEPseq2). In particular, the pipeline includes an additional step to analyze the added 3’ end U-tail. This additional script, coded in Python 3, extracts and analyzes the composition and the length of the sequence between the end of the detected poly(A) tail and the beginning of the 3’ adapter. This sequence was referred to as an additional tail. Additional tails containing at least 70% of Ts were considered as uridylated. All calculations and graphs of the present study were performed for annotated nuclear-encoded mRNAs, based on TAIR10. Falsely or ambiguously annotated RNAs were filtered out (see a list of filtered out gene identifiers in Supplementary Data Set S8). Bulk uridylation percentages were calculated using the total number of sequences as denominator. For per-transcript uridylation percentages, uridylation percentages were calculated for all mRNAs detected with at least 50 reads in each replicate and boxplots show the distribution of these percentages across mRNAs for each sample and condition. To plot the distribution of poly(A) tail sizes in Figure 3A and in Figure 7A, percentages of reads according to poly(A) tail size (1-nt bin) were calculated using the total number of sequences as denominator. To plot the per-transcript distribution of poly(A) tail sizes in Figures 7C and Supplemental Figure S5C, percentages of reads according to poly(A) tail size (2-nt bin) were calculated for each mRNA detected with at least 50 reads in each replicate and poly(A) profiles show the median (line), the first and the third quartile (grey shadow) of the percentages across mRNAs. For heatmaps, percentages of uridylation and percentages of reads according to poly(A) tail sizes (5nt-bin) in Supplementary Figure S1 and Figure 4, respectively, were calculated for each mRNA detected with at least 50 reads in each replicate and tissues and were z-score rescaled. The degradation pattern analysis (Figure 8) were based on the “end_polyA_type” tag defined by the original FLEP-seq2 pipeline (Jia et al. 2022). A read that doesn’t overlap the first exon or the last exon was considered as 5’- or 3’- fragmented, respectively.

Source data for figures are available in Supplementary Data Set S2. Statistical analysis results are available in Supplementary Data Set S9 and a table combining FLEP-seq2 and RNAseq data in dry seeds is provided in Supplementary Data Set S10. FLEP-seq2 results are also available in RNAvis, a web application developed in our institute using Rshiny, accessible at https://nebula.ibmp.unistra.fr/RNAvis/. Finally, mapping files were deposited to the Sequence Read Archive and are accessible through accession number PRJNA1286755.

### Analysis of synthetic DNA fragments harboring 0 to 6 Ts

Synthetic DNA fragments harboring a 10-nt poly(A) tail followed by 0, 1, 2, 3 or 6 Ts were synthesized as gBlocks Gene Fragments (Integrated DNA Technologies, Supplementary Data Set S7). gBlocks Gene Fragments were cloned in pGEM-T Easy (Promega) using the TA cloning method according to manufacturer’s instructions. Plasmids were purified using the Nucleospin Plasmid EasyPure kit (Macherey-Nagel) and their sequences verified by Sanger sequencing. Each validated cloned DNA fragment was then PCR-amplified from 10 ng of purified plasmid for 14 cycles using the PrimeSTAR GXL DNA polymerase (TaKaRa, R050A) and barcoded primers from the PCR-cDNA Barcoding kit (ONT, SQK-PCB111.24) as described for FLEP-seq2 library preparation. 1.25 fmol of each library were then sequenced alongside other samples on a P2 PromethION 2 Solo for ∼72 hours (Flow Cell FLO-PRO002, R9.4.1). Computational analysis of these synthetic DNA fragments was performed as described for the FLEP-seq2 library using the sequence of synthetic DNA fragments as the reference for mapping. Source data for figures are available in Supplementary Data Set S1.

### Uridylation analysis of FLEP-seq2 data from (Jia et al. 2022)

Uridylation analysis was performed from mapping .bam files available on China National Center for Bioinformation with accession OMIX881 using a homemade Python script (Extract_unmappedseq_final3.py), available on Github (https://github.com/jackson-peter/FLEPseq2) The script extracts the unmapped sequence found at the 3’ end of RNAs from the soft-clipped sequences of the mapping files, then searches for the nanopore adapter sequence and finally extracts the poly(A) tail and the eventual other added nucleotides. Additional tails containing at least 70% of Ts were considered as uridylated. As previously, all calculations were then performed for annotated nuclear-encoded mRNAs, based on TAIR10 and ambiguously annotated RNAs were filtered out (see a list of filtered out gene identifiers in Supplementary Data Set S8). Source data for figures are available in Supplementary Data Set S1. Statistical analysis results are available in Supplementary Data Set S9.

### Preparation of RNA-seq libraries

RNAseq libraries were generated from three biological replicates of flowers and after-ripened seeds using the same RNAs as those analyzed by FLEP-seq2. Total RNA was treated with DNase I (Thermo Fisher Scientific, EN0521) and purified using the RNeasy MinElute Clean-up (Qiagen, 74204). RNA-seq libraries were then produced from 500 ng of DNase-treated RNAs using the TruSeq Stranded Total RNA Library Prep Plant (Illumina, 20020610) according to manufacturer’s instructions. Sequencing was performed on an Illumina NextSeq 2000 system by the GenomEast platform (paired-end sequencing, 2 × 50 bases).

### Computational analysis for RNAseq

Reads have been mapped against the TAIR10 genome using hisat2 (v.2.2.1), with the following options: -p 4 -k 20 --max-intronlen 2000 --rna-strandness R. Read counts were then extracted for each representative transcript using FeatureCounts (v.2.0.6) and the Araport11 annotation (“Araport11_GTF_genes_transposons.Jul2023.gtf”) by specifying “gene” as feature type (-t option) and “gene_id” as attribute type (-g option). Differential expression analysis was performed in R (v.4.2.2) using the DESeq2 package (Love et al. 2014) (v1.38.3).

### RT-qPCR analysis for *DOG1* mRNA

For RT-qPCR analyses, RNA was extracted from seeds harvested from four independent plants using a phenol–chloroform-based extraction protocol. Seeds were ground to a fine powder while frozen and mixed with RNA extraction buffer (100 mM Tris pH 8.0, 5 mM EDTA,100 mM NaCl, 0.5% SDS, 1% beta-mercaptoethanol). RNA was extracted through sequential phenol–chloroform extractions, each followed by centrifugation at 14,000 g at 4°C. The RNA-containing supernatant was precipitated with 3M sodium acetate (pH 5.2) and isopropanol, then pelleted, washed, and resuspended in Milli-Q water. DNAse treatment of RNA was performed following the TURBO DNase protocol (ThermoFisher). RNA quality was assessed using agarose gel electrophoresis, Nanodrop 2000 spectrophotometer and PCR to check the absence of genomic DNA contamination. Reverse transcription was performed with SuperScript III (Invitrogen) according to the manufacturer’s protocol using a mixture (1:1) of random hexamers and oligo(dT). qPCR was performed using a LightCycler 480 real-time system (Roche) with SYBR Green mix with primers listed in Supplementary Data Set S7. Ct values of technical replicates were averaged for each genotype and biological replicates. Ct values were then normalized against the expression level of the housekeeping gene *UBC21* (AT5G25760) (Czechowski et al. 2005) by subtracting the Ct value of the *UBC21* control from the Ct value of *DOG1* for each sample. Obtained delta Ct (dCT) values were subsequently transformed using the 2^-dCt^ formula.

### Statistical analysis

All statistical analysis were performed were using R (v. 4.2.2) on RStudio (v.2023.12.1+402). Global percentages (Figure 1A and C, Figure 2B and D, Figure 3B, Figure 5A and B, Figure 7A, Figure 8C, Figure 9A-C, and Supplementary Figure S6B), were statistically compared using the R packages stats (v4.2.2) and car (v3.1-2) applying a generalized linear model for proportions with a quasibinomial distribution. To test the difference between distributions (Figure 1B, Figure 2C, Figure 5A, Figure 7C, Figure 8A, B and D, Supplementary Figure S1C and F, Supplementary Figure S2A, Supplementary Figure S3A and Supplementary Figure S5C) medians were calculated for each replicate and samples and a linear regression (R package stats, v4.2.2) was applied, as the Shapiro-Wilk test did not show evidence of non-normality. To compare FLEP-seq2 data with genomic features in Figure 1D and in Supplementary Figure S2B, the statistical analysis was performed using Pairwise Wilcoxon Rank Sum Tests with data considered as unpaired (non-parametric test, two-tailed). The multcomp R package (1.4-26) with Tukey contrasts was used for multiple comparison post hoc tests and the calculation of adjusted p-values. Differential expression analysis of RNAseq data was performed using the DESeq2 package (Love et al. 2014) (v.1.38.3). For all multigroup analyses, p-value were adjusted (Benjamini-Hochberg). For all statistical analyses, a p-value or an adjusted p-value of 0.05 was defined as the threshold of significance except for the analysis performed in Figure 1D and Supplementary Figure S2B for which the p-value threshold was set at 0.0001, due to the high number of observations. For the analysis of *DOG1* expression by RT-qPCR (Figure 9E), a two-tailed Wilcoxon rank-sum test for unpaired data (non-parametric test, two-tailed) was used to compare normalized and transformed Ct value (2^-dCt^). Detailed results of the statistical analysis are provided in Supplementary Data Sets S3 and S4 (RNAseq analysis) and Supplementary Data Set S9 for all other tests.

### Other analyses

All plots were generated using R (v. 4.2.2) and the R package ggplot2 (v. 3.5.1) on RStudio (v.2023.12.1+402), except for heatmaps that were generated using the pheatmap R package (v. 1.0.12). Boxplots shown in the paper displays the median, first and third quartiles (lower and upper hinges), the largest value within 1.5 times the interquartile range above the upper hinge (upper whisker) and the smallest value within 1.5 times the interquartile range below the lower hinge (lower whiskers). Publicly available RNAseq data from (Klepikova et al. 2016) (http://travadb.org) were used to analyze gene expression during seed development (Figure 6B, Supplementary Figure S4C and Supplementary Figure S5B). For heatmap in Figure 6B and Figure S5B, normalized read count (TMM, EdgeR) of each mRNA were divided by the maximum value of expression level among development stages, so all values vary from 0 to 1. As a reference for seed development stage, expression of mRNAs shown to be expressed at specific seed stage were also analyzed using RNAseq data from (Klepikova et al. 2016, Supplementary Data Set S5). All gene ontology analyses performed in the present paper were performed using DAVID (Huang et al. 2009b, 2009a), using an adjusted p-value of 0.05 (FDR) as threshold of significance.

### Accession Numbers

RNAseq datasets generated during this study have been deposited in NCBI’s Gene Expression Omnibus (Edgar et al. 2002) and are accessible through the GEO Series accession number GSE284704. Mapping files (.bam) for the re-analysis of FLEP-seq2 data from (Jia et al. 2022) were downloaded from the China National Center for Bioinformation with accession OMIX881. Mapping files (.bam) for FLEP-seq2 data sets generated in the present study have been deposited in the Sequence Read Archive and are accessible through accession number PRJNA1286755. Sequence data for the genes mentioned in this article can be found in the Arabidopsis Information Resource (TAIR, https://www.arabidopsis.org/) under the following accession numbers: AT2G45620 (*URT1*), AT2G39740 (*HESO1*), *DOG1* (AT5G45830).

## List of supplementary data files

Supplementary Figure S1. FLEP-seq2 to analyze U-tailing in different Arabidopsis plant tissues. Supports Figure 1.

Supplementary Figure S2. Changes of U-tail profiles during seed development and germination. Supports Figure 2.

Supplementary Figure S3. Analysis of mRNAs differentially uridylated in *urt1-1*, *heso1-4* and *heso1-4 urt1-1*. Supports Figure 5.

Supplementary Figure S4. Expression of *URT1* and *HESO1* during seed development. Supports Figure 6.

Supplementary Figure S5. URT1 inactivation triggers a massive deadenylation of mRNAs associated to the seed maturation program. Supports Figure 7.

Supplementary Figure S6. Impact of URT1 inactivation on seed germination. Supports Figure 9.

Supplementary Data Set S1. Source data related to spike-in and uridylation analysis of FLEP-seq data from Jia et al. 2022.

Supplementary Data Set S2. Source data related to FLEP-seq data generated in the present study.

Supplementary Data Set S3. Source data related to RNAseq analysis in seeds. Supplementary Data Set S4. Source data related to RNAseq analysis in flowers.

Supplementary Data Set S5. Expression data from publicly available RNAseq data from Klepikova et al., 2016.

Supplementary Data Set S6. Source data related to germination assays and *DOG1* expression.

Supplementary Data Set S7. List of oligonucleotides and synthetic DNA fragment used in this study.

Supplementary Data Set S8. List of AGI filtered out of the FLEP-seq2 analysis.

Supplementary Data Set S9. Source data for statistical analyses.

Supplementary Data Set S10. Table combining FLEP-seq2 and RNA-seq data in dry seeds.

## Acknowledgments

This research was funded by Centre National de la Recherche Scientifique (CNRS) and by a research grant from the French National Research Agency ANR-22-CE12-0002. This work of the Interdisciplinary Thematic Institute IMCBio, as part of the ITI 2021-2028 program of the University of Strasbourg, CNRS and Inserm, was supported by IdEx Unistra (ANR-10-IDEX-0002), and by SFRI-STRAT’US project (ANR 20-SFRI-0012) and EUR IMCBio (ANR-17-EURE-0023) under the framework of the French Investments for the Future Program. Nanopore sequencing was performed at the IBMP Gene Expression Analysis Platform. The RNA-seq sequencing was performed by the GenomEast platform, a member of the “France Génomique” consortium (ANR-10-INBS-0009). The authors thank Abdelmalek Alioua and Sandrine Koechler from IBMP Gene Expression Analysis Platform for help in Nanopore sequencing and RNA-seq libraries preparation, respectively, and David Pfliegler of the IBMP bioinformatics core facility for help in bioinformatic analysis. The authors thank Anthony Gobert for critical reading.

## Author Contributions

H.Z. and D.G. acquired funding; H.Z. conceived and supervised the research; J.P. conceived and developed analytical pipelines; J.R. performed experiments with the assistance of S.Sa., E.U., B.L.; J.P. and H.Z. analyzed data; H.Z., J.P., J.R., S. Sa, S.Sw, D.G. discussed the data. H.Z. drafted the paper. D.G. J.P., J.R., S. Sa, S.Sw edited and revised the paper.

