## Supplementary Figures for "The TUTase URT1 regulates the transcriptome of seeds and their primary dormancy"

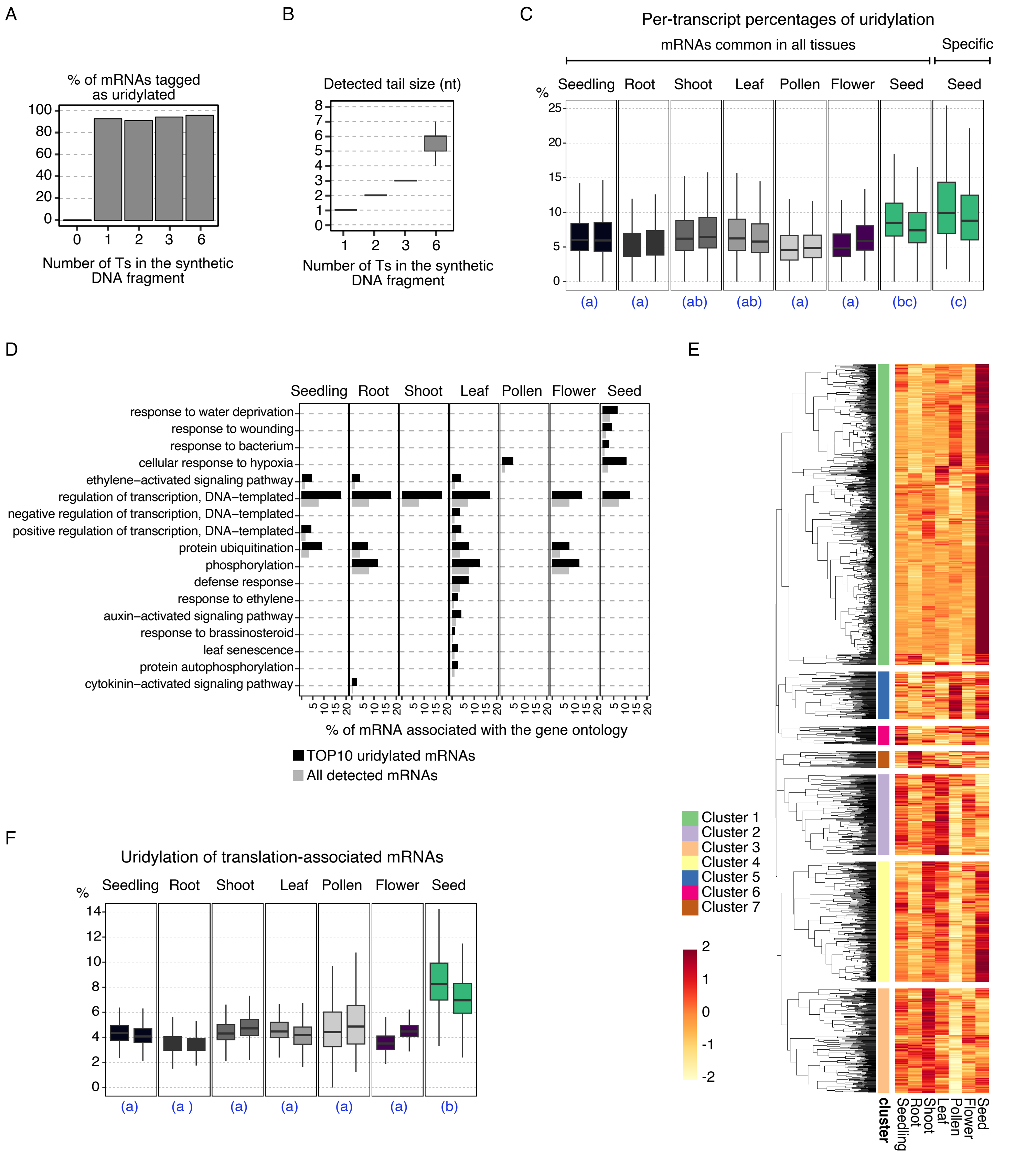


**Supplementary Figure S1: FLEP-seq2 to analyze U-tailing in different Arabidopsis plant tissues. Supports Figure 1.** A and B, Analysis of synthetic DNA fragments harboring a 10-nt poly(A) tail followed by 0 to 6 thymidines. The FLEP-seq2 pipeline from Jia et al. 2022 was adapted to analyze U-tail level and composition. A, Percentage of U-tail detected for each synthetic DNA fragment containing thymidines. B, U-tail sizes measured for each synthetic DNA fragment. C-F, U-tail analysis of FLEP-seq2 data from Jia et al. 2022. Data are shown for two replicates. C, Per-transcript percentages of uridylation for mRNAs expressed in all tissues or specifically expressed in seeds. Percentages were calculated for mRNAs detected with at least 50 reads for each replicate and tissue or for mRNAs specifically detected in seeds. D, Biological processes significantly enriched among the 10% most uridylated mRNAs (TOP10 uridylated mRNAs). Gene ontology analysis based on DAVID (https://david.ncifcrf.gov, adjusted p-value <0.05). Black and grey bar plots represent the proportion of mRNAs associated to each biological process significantly enriched among TOP10 uridylated or among all mRNAs, respectively. E, Heatmap showing the variation patterns of uridylation percentages (z-score rescaled) between tissues. Transcripts were clustered based on their similarity of uridylation profiles (k=7). F, Per-transcript percentages of uridylation for translation-associated mRNAs. Each boxplot in (B),(C) and (F) displays the median, first and third quartiles (lower and upper hinges), the largest value within 1.5 times the interquartile range above the upper hinge (upper whisker) and the smallest value within 1.5 times the interquartile range below the lower hinge (lower whiskers). In (C) and (F), significantly different values are labelled by different letters (linear model, adjusted p-value < 0.05, n=3).

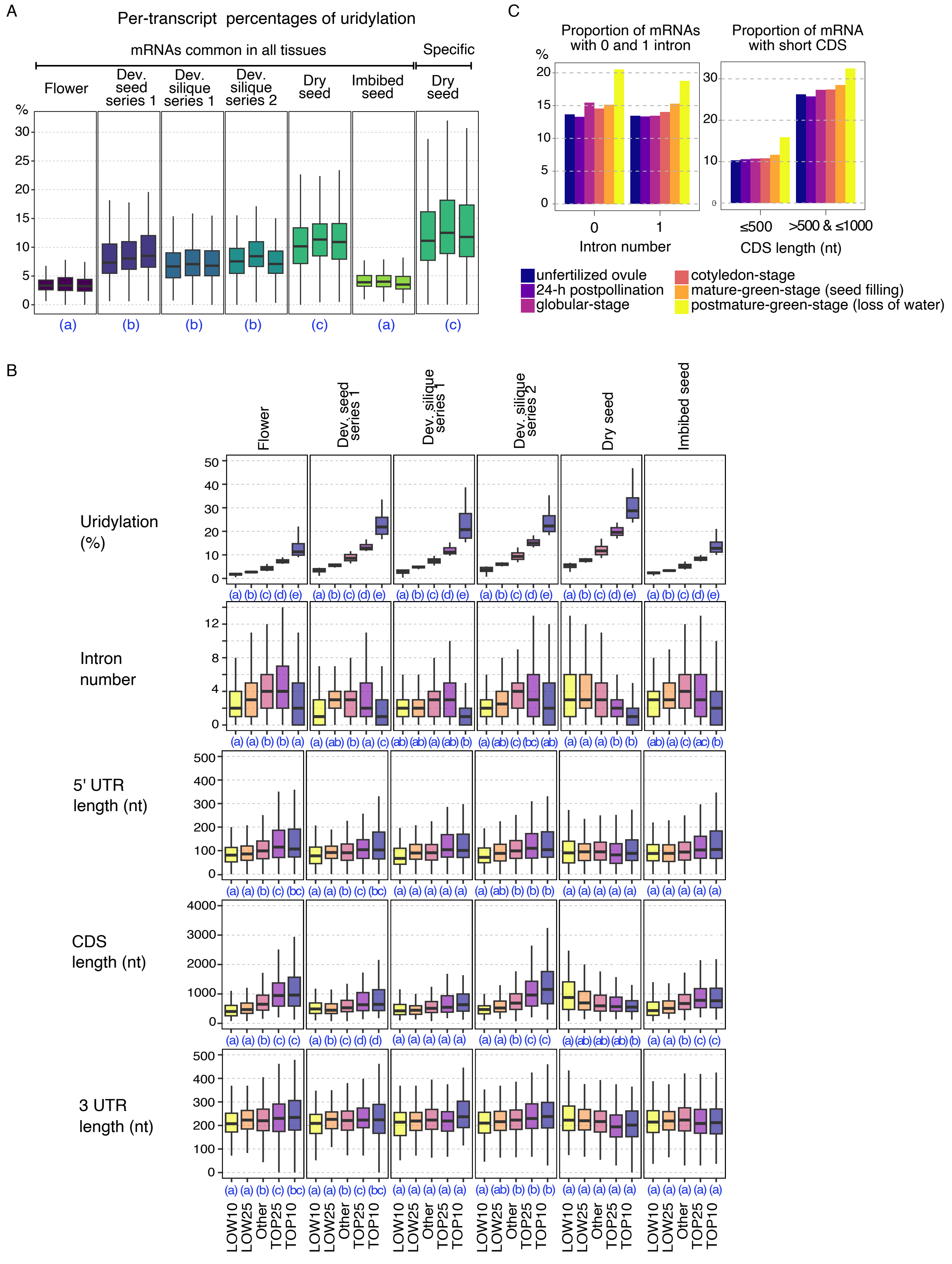


**Supplementary Figure S2. Changes of U-tail profiles during seed development and germination. Supports Figure 2.** FLEP-seq2 data from flowers, developing seeds and siliques (containing seeds), dry seeds and 24h-imbibed seeds. Data are shown for three replicates. A, Per-transcript percentages of uridylation for mRNAs expressed in all tissues or specifically expressed in seeds. Percentages were calculated for mRNAs detected with at least 50 reads for each replicate and tissue or for mRNAs specifically detected in dry seeds. Each boxplot displays the median, first and third quartiles (lower and upper hinges), the largest value within 1.5 times the interquartile range above the upper hinge (upper whisker) and the smallest value within 1.5 times the interquartile range below the lower hinge (lower whiskers). Significantly different values (linear model, adjusted p-value < 0.05, n=3) are labelled by different letters. B, Feature analysis for the 10% and 25% most and less uridylated mRNAs or for other mRNAs. Each boxplot displays the median, first and third quartiles (lower and upper hinges), the largest value within 1.5 times the interquartile range above the upper hinge (upper whisker) and the smallest value within 1.5 times the interquartile range below the lower hinge (lower whiskers). Significantly different values are labelled by different letters (adjusted p-value < 0.0001, unpaired and two-tailed pairwise Wilcoxon rank sum tests, n > 100). C, Proportion of mRNAs with a short CDS and a low number of introns among mRNA specifically expressed at different development stages (based on Le et al. 2010).

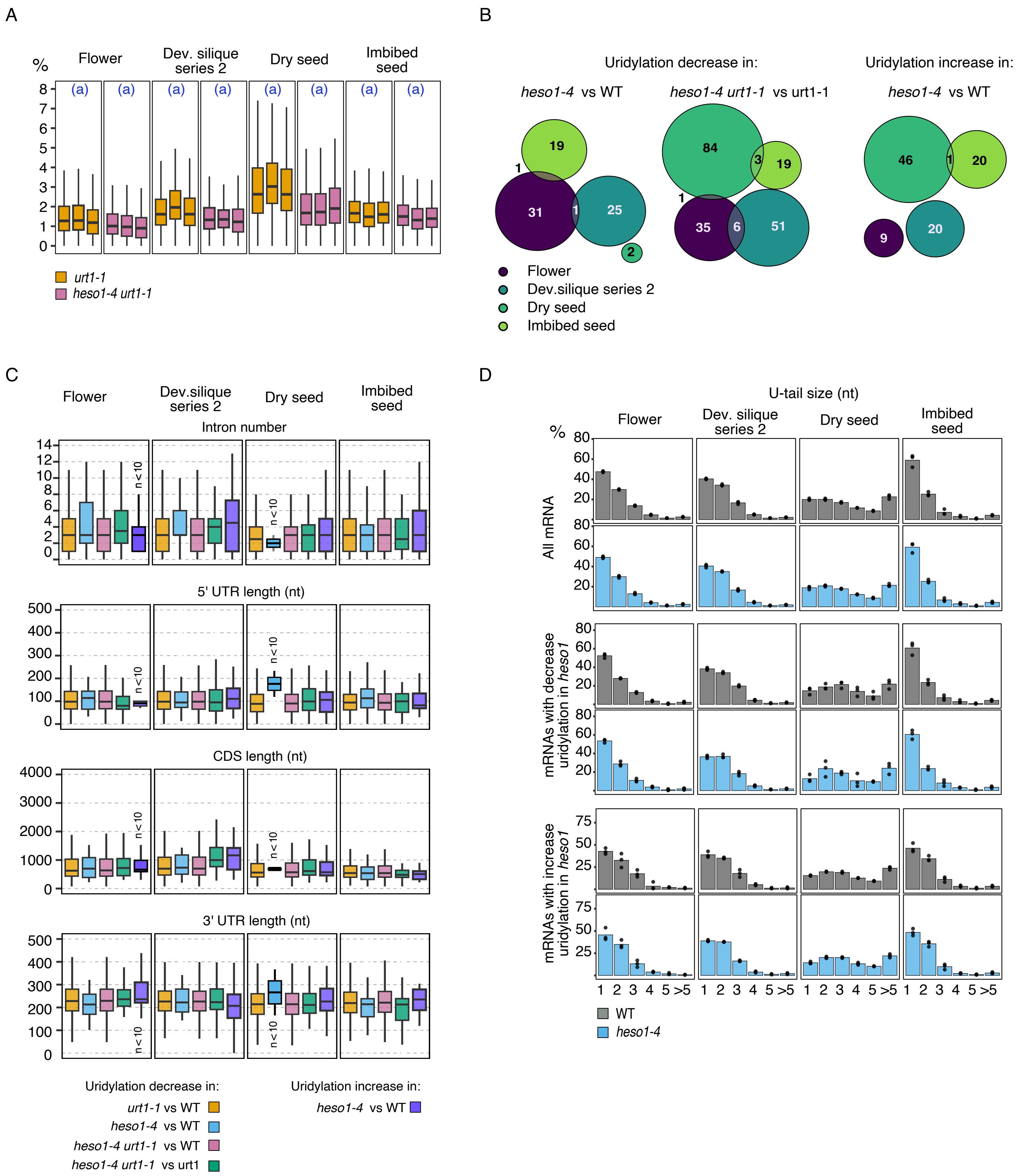


**Supplementary Figure S3. Analysis of mRNAs differentially uridylated in *urt1-1*, *heso1-4* and *heso1-4 urt1-1*. Supports Figure 5.** A, Per-transcript percentages of uridylation in *urt1-1* and *heso1-4 urt1-1*. Percentages were calculated for mRNAs detected with at least 50 reads for each replicate and genotype. Significantly different values are labelled by different letters (linear model, adjusted p-value < 0.05, n=3) . B, Venn diagrams showing the intersection between the different lists of mRNAs. C, Feature analysis for the different list of mRNAs. In (A) and (C), Boxplot displays the median, first and third quartiles (lower and upper hinges), the largest value within 1.5 times the interquartile range above the upper hinge (upper whisker) and the smallest value within 1.5 times the interquartile range below the lower hinge (lower whiskers). D, % of tails according to the number of Us in WT and in *heso1-4* for mRNAs showing a decrease or an increase uridylation percentage in *heso1-4*. Individual points and the bar plot show individual replicates and the average, respectively. Bar plots for *urt1-1* and *heso1-4 urt1-1* plants are not shown as uridylation is very low.

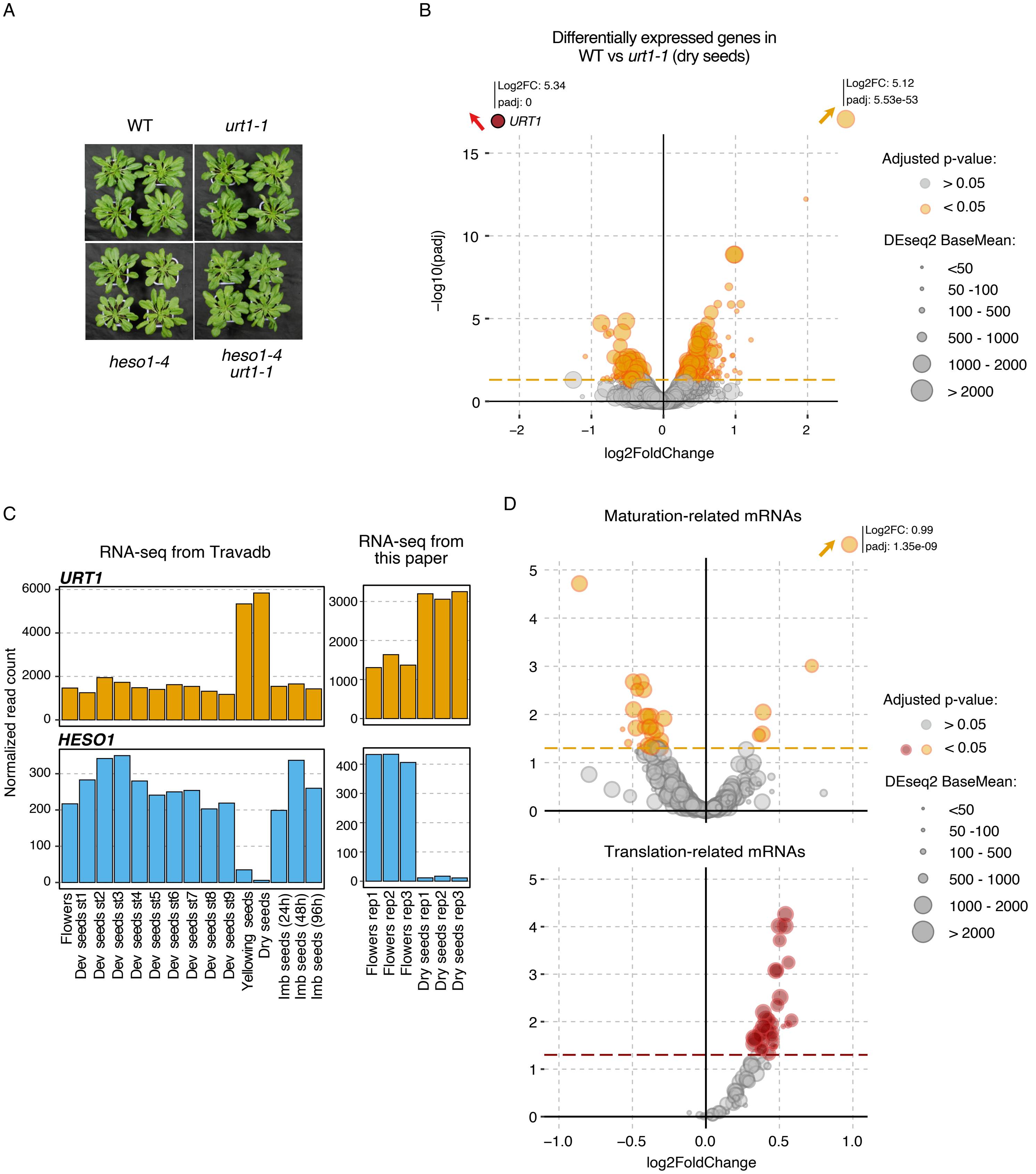


**Supplementary Figure S4. Impact of URT1 inactivation on seed transcriptome and expression of *URT1* and *HESO1* during seed development. Supports Figure 6.** A, Representative picture of eight- week-old plants for WT, *urt1-1*, *heso1-4* and *heso1-4 urt1-1*. B, Volcano plot showing transcripts significantly differentially accumulated in *urt1-1* mutant. C, Transcript accumulation of *URT1* and *HESO1* during seed development based on publicly available RNAseq data from Klepikova et al., 2016 (left panel, EdgeR TMM normalization) and on RNAseq data from this paper (right panel, DEseq2 normalization). D, Volcano plot showing transcripts significantly differentially accumulated in *urt1-1* mutant for maturation- and translation-related mRNAs. For volcano plots in (B) and (D), the dashed line indicates the significant threshold (adjusted p-value < 0.05, DEseq2, n=3). Dots represent individual mRNAs. Orange or red dots represent mRNAs with an adjusted p-value (padj) < 0.05. Grey dots show no significant difference. The point size was defined according to the expression level of each mRNA (BaseMean, DEseq2).

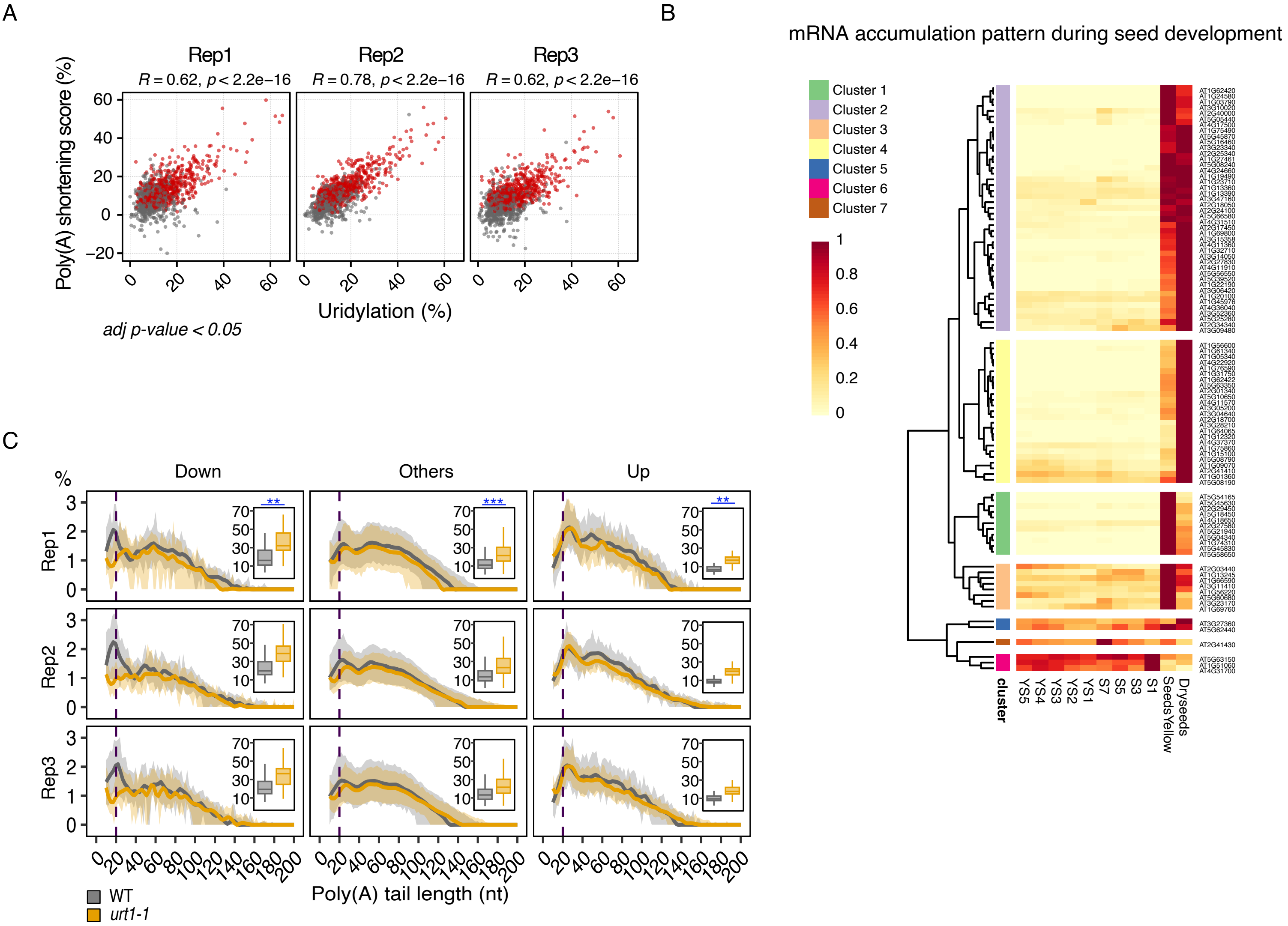


**Supplementary Figure S5. URT1 inactivation triggers a massive deadenylation of mRNAs associated to the seed maturation program. Supports Figure 7.** A, Poly(A) shortening scores according to uridylation percentages. Poly(A) shortening score corresponds to the difference of the percentage of A-tails shorter than 10As between *urt1-1* and WT dry seeds. Red points show mRNAs with a significant increase accumulation of $A_{<10}$-tails (adjusted p-value <0.05, generalized linear model for proportion with a quasibinomial distribution, n=3). Pearson coefficients of correlation are shown with their associated p-value (ggpubr R package). B, Heatmap showing the expression pattern during seed development of the 93 mRNAs that are the most impacted in term of poly(A) shortening. Samples correspond to different development stages of isolated seeds from early development stages (young seeds, YS and seeds, S) to desiccating seeds and dry seeds (see Figure 6 and Supplemental Table S5). Expression data are publicly available RNAseq data from Klepikova et al., 2016. For each mRNA, normalized read count (TMM, EdgeR) were divided by the maximum value of expression level among development stages, so all values vary from 0 to 1. Transcripts were clustered based on their similarity of expression profiles (k=7). C, Comparison of the poly(A) tail sizes between WT and *urt1-1* for mRNAs found to be down or up-regulated by *urt1-1* mutation. Distribution of poly(A) tail sizes from 10 to 200 As. Percentages were calculated using the total number of sequences as denominator, including those with A-tails < 10 As. The lines and the grey shadow show the median and the first and third quartiles of percentages across mRNAs, respectively. Boxplots next to each distribution show the per-transcript proportion of reads with no or short A-tail (<10As). Boxplot displays the median, first and third quartiles (lower and upper hinges), the largest value within 1.5 times the interquartile range above the upper hinge (upper whisker) and the smallest value within 1.5 times the interquartile range below the lower hinge (lower whiskers). Significantly different values (adjusted p-value < 0.05, linear model, n=3) are labelled by stars (* <0.05, *<0.01, ***<0.001).

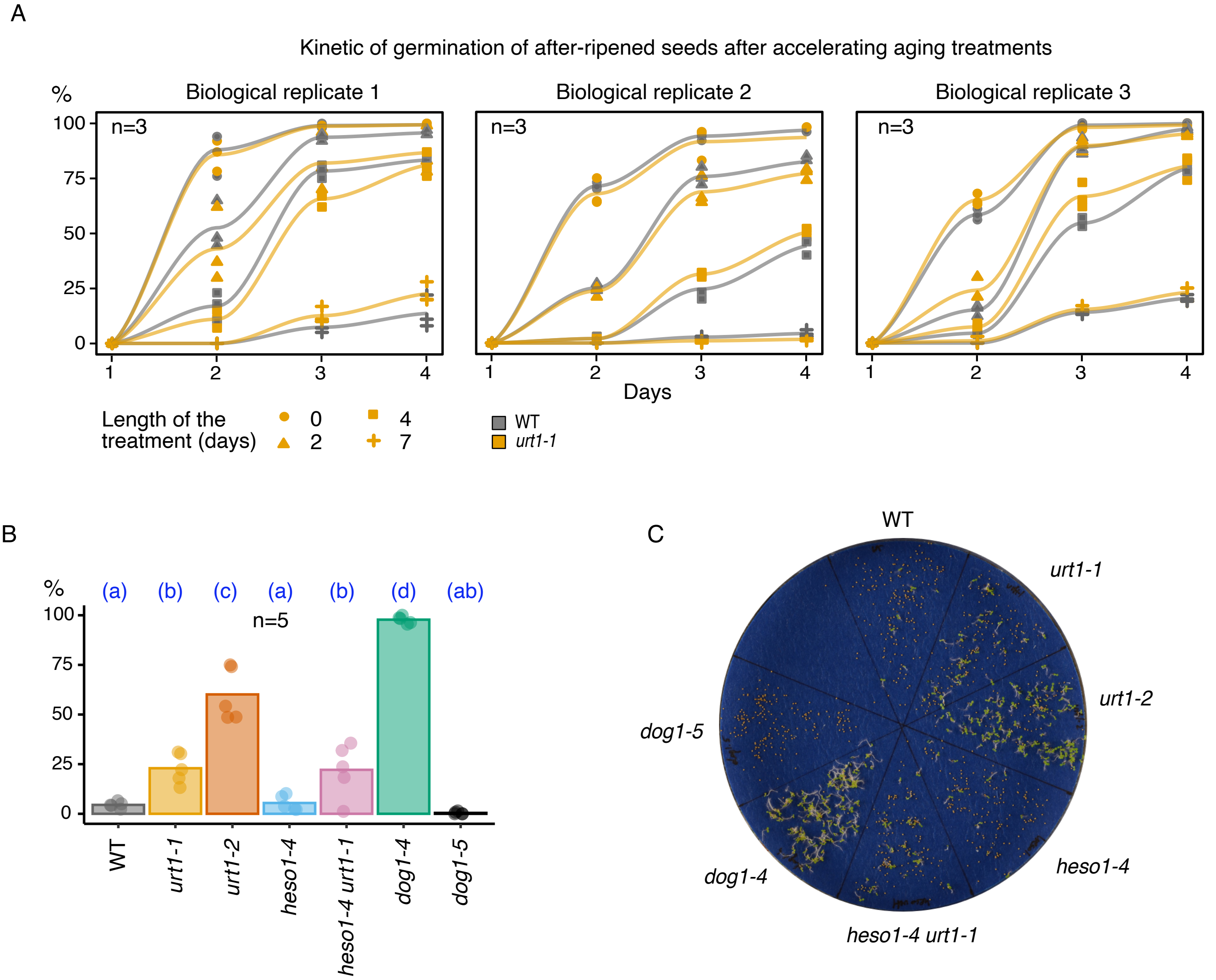


**Supplementary Figure S6. Impact of URT1 inactivation on seed germination. Supports Figure 9.** A, Kinetic of germination of WT and *urt1-1* after-ripened seeds for different durations of accelerated aging treatment. Germination assays were repeated for three biological replicates. Individual points show individual technical replicates, and line plots show the average. B, Level of primary dormancy of WT, *urt1-1*, *urt1-2, heso1-4*, *heso1-4 urt1-1* freshly harvested seeds. *dog1-4* and *dog1-5* were included as positive and negative controls, respectively. In B and C, the bar plots show the percentage of germinated seeds after four days. Individual points show the percentages for seeds harvested from five individual plants and bar plots show the average. C, A representative image of germination assay for freshly harvested seeds is shown. In all graphs, significantly different values are labelled by different letters (adjusted p-value < 0.05, generalized linear model for proportion, quasibinomial distribution, n is indicated on each graph).
